# Seasonal and spatial transitions in phytoplankton assemblages spanning estuarine to open ocean waters of the tropical Pacific

**DOI:** 10.1101/2024.05.23.595464

**Authors:** Sarah J. Tucker, Yoshimi M. Rii, Kelle C. Freel, Keliʻiahonui Kotubetey, A. Hiʻilei Kawelo, Kawika B. Winter, Michael S. Rappé

## Abstract

Islands in the tropical Pacific supply elevated nutrients to nearshore waters that enhance phytoplankton biomass and create hotspots of productivity in otherwise nutrient-poor oceans. Despite the importance of these hotspots in supporting nearshore food webs, the spatial and temporal variability of phytoplankton enhancement and changes in the underlying phytoplankton communities across nearshore to open ocean systems remain poorly understood. In this study, a combination of flow cytometry, pigment analyses, 16S rRNA gene amplicons, and metagenomic sequencing provide a synoptic view of phytoplankton dynamics over a four-year, near-monthly time-series across coastal Kāneʻohe Bay, Hawaiʻi, spanning from an estuarine Indigenous aquaculture system to the adjacent offshore environment. Through comparisons with measurements taken at Station ALOHA located in the oligotrophic North Pacific Subtropical Gyre, we observed a sharp and persistent transition between picocyanobacterial communities, from *Synechococcus* clade II abundant in the nearshore to *Prochlorococcus* HLII proliferating in offshore and open ocean waters. In comparison to immediately adjacent offshore waters and the surrounding open ocean, phytoplankton biomass within Kāneʻohe Bay was dramatically elevated. Members of the phytoplankton community revealed strong seasonal patterns, while nearshore phytoplankton biomass positively correlated with wind speed, rainfall, and wind direction, and not water temperatures. These findings elucidate the spatiotemporal dynamics underlying transitions in ocean biogeochemistry and phytoplankton dynamics across estuarine to open ocean waters in the tropical Pacific and provide a foundation for quantifying deviations from baseline conditions due to ongoing climate change.

## Introduction

In marine ecosystems, phytoplankton play a crucial role by forming the base of the aquatic food web, where their productivity, abundance, cell size, and community composition are greatly influenced by light and nutrient availability (Azam et al. 1983; Cloern 1996; Acevedo-Trejos et al. 2015; Dutkiewicz et al. 2020). Surrounded by oligotrophic, open ocean waters, the coastal waters of remote islands in the tropical Pacific harbor a sharp increase in nutrients through physical, biological, geological, and anthropogenic processes that result in increased phytoplankton biomass, cell size, and productivity (i.e., the Island Mass Effect, or IME; Doty and Oguri 1956; Gove et al. 2016). The enhanced primary productivity boosts marine biodiversity (Messié et al. 2022), promotes carbon sequestration (Falkowski et al. 1998; Stukel et al. 2023), and increases secondary productivity, which in turn supports regional fisheries and other resources relied upon by island communities (Williams et al. 2015; Stock et al. 2017).

Elevated phytoplankton biomass from near-island coastal waters is important to maintaining healthy and productive coastal food webs, however these food webs may be vulnerable due to their close proximity to open ocean oligotrophic waters. One result of increasing sea surface temperatures from ongoing global climate change is that it has the potential to increase the intensity of water column stratification of the open ocean, which can trap nutrients at depths below where phytoplankton at the ocean’s surface can access them (Yamaguchi and Suga 2019; Li et al. 2020). This is expected to lead to a number of potential ecosystem impacts: an expansion of nutrient-poor “ocean deserts” in the open ocean gyres (Irwin and Oliver 2009; Fu et al. 2016), a shift towards small phytoplankton classes (Flombaum et al. 2020; Flombaum and Martiny 2021; Henson et al. 2021), a decline in global phytoplankton biomass and primary productivity (Boyce et al. 2010; Bopp et al. 2013), and an amplification of biomass declines across the trophic food web (Kwiatkowski et al. 2018; Moore et al. 2018). However, recent studies suggest that genetic and physiological adaptations of phytoplankton may sustain primary productivity even under the lower nutrient conditions that may occur with increased ocean stratification (Kwon et al. 2022; Martiny et al. 2022; Hagstrom et al. 2024).

An improved understanding of the spatiotemporal variability of biogeochemical conditions and phytoplankton assemblages across coastal to open ocean systems adjacent to island masses in the tropical Pacific could help inform management of local marine environments and larger ecosystem models. Nutrients are delivered to near-island waters by both land- (e.g., stream discharge, groundwater, anthropogenic impacts; Laws and Allen 1996; Drupp et al. 2011; McKenzie et al. 2019) and ocean-based processes (e.g. upwelling, internal waves; Gove et al. 2016; Friedrich et al. 2021). The exchanges of nutrients and phytoplankton across these adjacent environments have important implications for enhancing phytoplankton productivity (Messié et al. 2020; Leng et al. 2023) and structuring of community composition (Ribalet et al. 2010; Tucker et al. 2021; Comstock et al. 2022; Comfort et al. 2024). However, most studies examining phytoplankton dynamics in the tropical Pacific have sampled the offshore or the coastal environment in isolation (Pasulka et al. 2013; Estrada et al. 2016; Rii et al. 2016; Selph et al. 2018) or sampled both environments but typically with limited taxonomic information or temporal coverage (James et al. 2020; Vollbrecht et al. 2021; Messié et al. 2022). Coastal-ocean gradients in biogeochemistry and phytoplankton communities are often modified by seasonal or episodic changes in the environment (Cox et al. 2006; Haber et al. 2022; James et al. 2022). On high islands in the tropical Pacific, storm events can be particularly impactful with high stream discharge carrying a substantial proportion of the annual nutrient budget for coastal waters and invoking rapid phytoplankton responses (Carlo et al. 2007; Hoover and Mackenzie 2009). Furthermore, submarine groundwater discharge can contribute greater nutrient inputs than surface run-off (Dulai et al. 2016) and future climate change scenarios are estimated to substantially impact groundwater contributions to coastal systems (Ghazal et al. 2023).

To illuminate the factors influencing phytoplankton communities across near-island to open ocean environments in the tropical Pacific, this study examined the effect of spatial and temporal variability in biogeochemical conditions on phytoplankton communities across multiple habitats that link the coastal environment of Oʻahu, Hawaiʻi, with the offshore. These habitats span a tidally-influenced, estuarine environment within an Indigenous aquaculture system, through the interior of coastal Kāneʻohe Bay, and to the offshore ocean environment surrounding Kāneʻohe Bay. We also made comparisons to data collected by the Hawaii Ocean Time-series (HOT), a 30+ year time-series initiative measuring temporal trends of the adjacent oligotrophic North Pacific Subtropical Gyre (NPSG; Karl and Lukas 1996; Karl and Church 2014). Together, this extensive spatial and temporal coverage revealed nearshore enhancement of phytoplankton biomass, pronounced seasonality in nearshore biogeochemistry and phytoplankton biomass and composition, and distinct transitions in phytoplankton communities spanning <6 km to >100 km across Kāneʻohe Bay to the NPSG.

## Materials and Methods

### Study location

The Hawaiian archipelago within the oligotrophic NPSG is the world’s most remote island chain. Kāneʻohe Bay, located on the windward side of the island of Oʻahu (21° 28′ N, 157° 48′ W), is a well-studied, coral-reef-dominated embayment (**Fig. 1a-b**). The bay has a total surface area of 41.4 km^2^ and is approximately 4.3 km wide, 12.8 km in length, and 10 m deep on average (Jokiel 1991). Sharp nearshore to offshore gradients in biogeochemical parameters occur over a short distance (<6 km) along with a diverse topography due to patch, fringing, and barrier reefs (Jokiel 1991; Tucker et al. 2021). Localized freshwater input from streams contribute to episodic spatial variability in environmental conditions, including salinity and inorganic nutrient concentrations (Cox et al. 2006; Yeo et al. 2013; Tucker et al. 2021), with non-trivial contributions in inorganic nitrogen and silica via submarine groundwater discharge (Dulai et al. 2016). Water residence time within the bay varies from less than a day to over one month (Lowe et al. 2009), with the highest residence times in the sheltered southern lobe. Oceanic water, primarily driven by wave action, flows into the bay over a large barrier reef located in the central bay (Lowe et al. 2009). Water is generally transported out of the bay through two nearly-parallel channels positioned in the southern and northern portions of the bay. For most of the year, the bay is well mixed by northeasterly tradewinds. However, periods of high temperatures and low wind speeds can cause vertical stratification in the water column (Smith 1981).

**Fig. 1.**
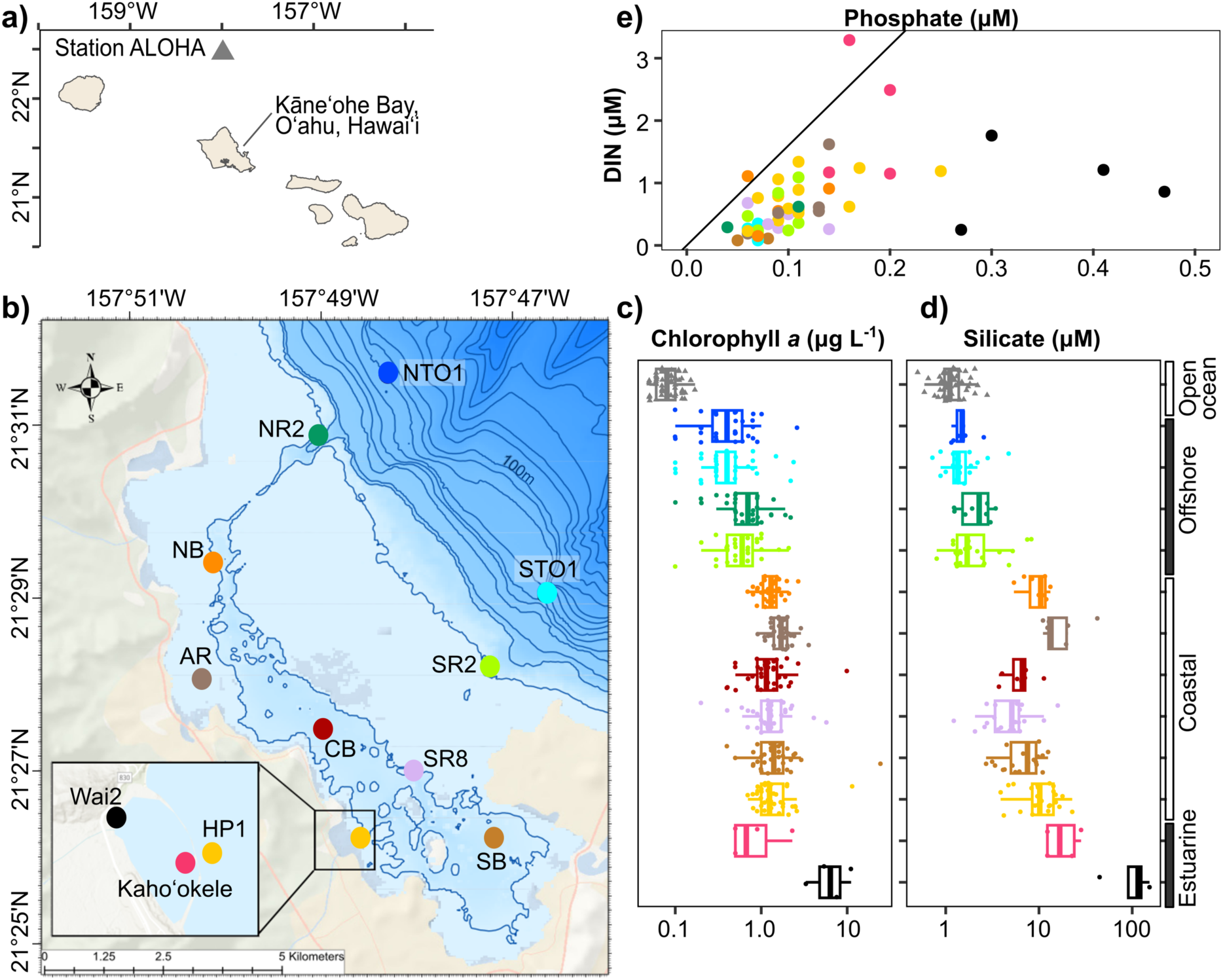
a) Location of Kāneʻohe Bay on Oʻahu, Hawaiʻi and the position of Station ALOHA (22° 45’N, 158°W), a sampling station of the Hawaii Ocean Time-series (HOT) program. b) Map of sampling stations located on the windward side of the island of Oʻahu Hawaiʻi. Inset shows two stations within Heʻeia Fishpond (Wai2 and Kahoʻokele) and one immediately adjacent to the fishpond (HP1). Contour lines mark every ten meters up until 100 m and then 50 m intervals for depths >100 m. c) Fluorometric chlorophyll *a* (Chla) and d) silicate concentrations (both plotted on a log scale) from the estuarine to open ocean environments examined in this study. e) Ratios of dissolved inorganic nitrogen (DIN):Phosphate. Diagonal line denotes Redfield ratio of 16:1 suggesting nitrogen limitation is characteristic of the system.

At the mouth of Heʻeia Stream in the southern section of Kāneʻohe Bay is an ∼800-year-old, 0.356 km^2^ Indigenous aquaculture system known contemporarily as Heʻeia Fishpond, but anciently as Pihi Loko Iʻa (Kelly 1973). Indigenous aquaculture systems in Hawaiʻi engaged in trophic engineering to promote primary productivity that sustained the population through abundant food fish and reef fish (Winter et al. 2020a). A 2.5 km basalt rock wall (kuapā) filled with coral rubble encompasses Heʻeia Fishpond. The wall is equipped with multiple sluice gates (mākāhā) that increase water residence time while still allowing for exchange between coastal Kāneʻohe Bay and Heʻeia stream waters. The mākāhā system size selects for juvenile fish exchange within the fishpond and Kāneʻohe Bay, allowing them to flourish within the high nutrient environment of the fishpond, while protecting them from predators. While it has been understood that the fishpond provided the perfect nursery habitat for prized food fish, little is known about the exchange of biogeochemically productive water and organisms from within the fishpond to the rest of Kāneʻohe Bay, and beyond.

### Collaboratively developed research

In following the collaborative research methodology practiced in the Heʻeia National Estuarine Research Reserve (NERR; Winter et al. 2020b; Kūlana Noiʻi Working Group 2021), and guided by interests expressed and knowledge possessed by the Indigenous (*Kānaka ʻŌiwi*, Native Hawaiian) stewards of Heʻeia Fishpond, we sought to establish a current baseline understanding of phytoplankton biomass and community composition from within the estuarine fishpond environment out to open ocean waters adjacent to Oʻahu. All individuals who developed the methods also collaborated on interpreting the data and are listed as co-authors on this paper. Sampling campaigns were conducted with permission from Paepae o Heʻeia, the organization that stewards Heʻeia Fishpond, and the private landowner, Kamehameha Schools and Bishop Estate.

### Sample collection and environmental parameters

Between August 2017 and June 2021, seawater was collected from a depth of 2 m at 10 sites in Kāneʻohe Bay and the adjacent offshore waters on a near-monthly basis (36 sampling events over 46 months) as part of the Kāneʻohe Bay Time-series (KByT) using previously described methods (**Fig. 1b, Supporting Information Table S1**; Tucker et al. 2021). Between September 2020 and June 2021, two additional stations within Heʻeia Fishpond were also sampled at a quarterly interval **(Fig. 1b**): Station Wai2 (now known as Waimāʻama), located in the northwestern corner of the Fishpond within the Heʻeia Stream mouth (known as muliwai) is a highly turbid and brackish water environment with tidal fluctuations resulting in salinity ranges of 10-35 ppt (Wai2, NERR Centralized Data Management Office: https://cdmo.baruch.sc.edu/).

Station Kahoʻokele (Kaho) is located at a sluice gate facing the ocean and receives high exchange with coastal Kāneʻohe Bay (∼25-35 ppt, Möhlenkamp et al. 2019; NERR Centralized Data Management Office: https://cdmo.baruch.sc.edu/). At all stations, seawater samples for biogeochemical analyses and nucleic acids were collected, as were *in situ* measurements of seawater temperature, pH, and salinity with a YSI 6600 or ProDSS multi-parameter sonde (YSI Incorporated, Yellow Springs, OH, USA). Approximately one liter of seawater was prefiltered with 85-μm Nitex mesh and subsequently filtered through a 25-mm diameter, 0.1-μm pore-sized polyethersulfone (PES) filter membrane (Supor-100, Pall Gelman Inc., Ann Arbor, MI, USA) to collect microbial cells for DNA isolation. The filters were subsequently submerged in DNA lysis buffer (Suzuki et al. 2001; Yeo et al. 2013) and stored in −80°C until further processing.

Seawater subsamples for fluorometric chlorophyll *a* concentrations (125 mL) and photosynthetic pigments via high-performance liquid chromatography (HPLC; 2 L) were collected on 25-mm diameter GF/F glass microfiber filters (Whatman, GE Healthcare Life Sciences, Chicago, IL, USA) and stored for up to 24 months in aluminum foil at −80°C until extraction. The collection of phytoplankton pigments on the GF/F glass microfiber filters allow for comparisons with the HOT data. However, because the filters have a pore size of 0.7 µm, we acknowledge that most small cyanobacteria were likely missed. Chlorophyll *a* was extracted with 100% acetone and measured with a Turner 10-AU fluorometer (Turner Designs, Sunnyvale, CA, USA) following standard techniques (Welschmeyer 1994). Photosynthetic pigments measured via HPLC were extracted in 100% acetone and analyzed on a Waters 2690 separations module equipped with a C18 column and full spectrum photodiode array detector, following Mantoura and Llewellyn (1983) and modified according to Bidigare et al. (1989). Chlorophyll *a* concentrations measured via the fluorometer are herein referred to as chlorophyll *a* (Chla), while chlorophyll *a* concentrations measured by high performance liquid chromatography are specified as total chlorophyll *a* (Tchla), totaling monovinyl and divinyl chlorophyll *a*.

For cellular enumeration, seawater was preserved in 2 mL aliquots in a final concentration of 0.95% (v:v) paraformaldehyde (Electron Microscopy Services, Hatfield, PA, USA) at −80 °C until analyzed via flow cytometry. Cellular enumeration of cyanobacterial picophytoplankton (*Synechococcus* and *Prochlorococcus*), photosynthetic picoeukaryotes, and non-cyanobacterial (presumably heterotrophic) bacteria and archaea (hereafter referred to as heterotrophic bacteria) was performed on a Beckman Coulter CytoFLEX S, following the method of (Monger and Landry 1993). Inorganic nutrients were measured using a Seal Analytical AA3 HR Nutrient Autoanalyzer (detection limits: NO_2_^−^ + NO_3_^−^, 0.009 µM; SiO_4_, 0.09 µM; PO_4_^3 –^, 0.009 µM; NH_4_, 0.03 µM).

Rainfall, wind speed, and wind direction were monitored using data collected at a meteorological station located at the Hawai‘i Institute of Marine Biology (HIMB) on Moku o Lo‘e in Kāneʻohe Bay (http://www.pacioos.hawaii.edu/weather/obs-mokuoloe/). Either maximum (e.g., rainfall, wind speed) or average (wind direction) values were taken across 1- to 7-day windows leading up to the sampling event, depending on data availability from the station. One sampling event (February 5, 2021) had no data for the 7 days prior to sampling and so the data for a 30-day window were used. Metadata from Station ALOHA (∼5 m depth, August 2017 to December 2020 were downloaded from https://hahana.soest.hawaii.edu/hot/hot-dogs/, accessed on 9/12/2022).

Spatiotemporal comparisons of environmental variables, cellular abundances, and phytoplankton pigments were conducted using the R package ʻmultcomp’ (Hothorn et al. 2008) with one-way ANOVA testing for multiple comparisons of means with Holm correction and Tukey contrasts. Summer (28 June through 28 September) and winter (27 December through 29 March) seasons were defined using harmonic regression analyses of surface seawater temperature collected hourly between 2010–2019 at NOAA station MOKH1 in Kāneʻohe Bay (https://www.ndbc.noaa.gov/station_page.php?station=mokh1; Tucker et al. 2021).

Spatiotemporal variation in cellular abundances and phytoplankton pigments were visualized using ‘mba.surf’ from MBA (Finley et al. 2017) to interpolate data over the KByT sampling events and stations.

The map of Kāneʻohe Bay was plotted in ArcGIS Pro v2.9. Contour lines were drawn using bathymetry metadata (http://www.soest.hawaii.edu/hmrg/multibeam/bathymetry.php). The map of Oʻahu was plotted in R v 4.4.1 (R Core Team 2021) with ggplot2 v3.4.3 (Wickham 2016), using a shape file from the Hawaii Statewide GIS Program (https://prod-histategis.opendata.arcgis.com/maps/HiStateGIS::coastline).

### DNA extraction, 16S rRNA gene amplicon sequencing, & metagenome sequencing

DNA extraction (Qiagen Blood and Tissue Kit, Qiagen Inc., Valencia, CA, USA) and 16S rRNA gene sequencing followed previously published methods (Tucker et al. 2021). Briefly, amplicon libraries were made from polymerase chain reactions of the 16S rRNA gene using barcoded 515F and 926R universal primers complete with Illumina sequencing adapters, barcode and index (Parada et al. 2016) and paired-end sequencing with MiSeq v2 2x250 technology (Illumina, San Diego, CA, USA). Genomic DNA from 32 of the 368 total samples collected between 2017-2021 were used for metagenomic sequencing. This included samples from four sampling events between 2017 and 2019 at 6 to10 stations. Libraries were constructed from approximately 100 ng of genomic DNA using the Kappa HyperPrep Kit (Roche, Pleasanton, CA, USA) with mechanical shearing (Covaris, Woburn, MA, USA) and paired-end sequenced on a single lane of the NovaSeq 6000 SP 150 (Illumina, San Diego, CA, USA).

### Sequence analysis

Amplicon sequence data generated from KByT sampling between July 2019 and June 2021 were analyzed in conjunction with previously published amplicon data spanning August 2017 to June 2019 (PRJNA706753; Tucker et al. 2021). For each of the two sequencing runs, samples were demultiplexed and quality controlled using Qiime2 v2.4 (Bolyen et al. 2019). Full length forward reads (251 base pairs) were denoised and chimeras removed using DADA2 (Callahan et al. 2019) to delineate amplicon sequencing variants (ASVs). Reverse reads were not used because of inconsistent quality. ASVs were assigned taxonomy using SILVA v138 as a reference database (Quast et al. 2012) and the two runs were subsequently merged in Qiime2 using ʻfeature-table merge-seqs’ to yield a set of unique ASV sequences found across the two sequencing runs. ASVs that contained at least 10 reads in at least two samples were retained.

ASVs classified by the SILVA v138 database as Eukaryota, unassigned at the domain level, or classified as chloroplast at the order level were re-classified using the PR2 v 4.14.0 database (Guillou et al. 2013) in the DECIPHER v 2.20.0 R package (Wright 2016) using a 60% confidence threshold cut off. Sequences classified as Bacteria, Archaea, or chloroplast at the order-level (n= 4,117 ASVs) were retained for further analyses, while those unclassified at the domain level (n=100 ASVs) or classified as Eukaryota (18S rRNA sequences, n=337 ASVs) were excluded from further analyses. Eukaryota 18S rRNA sequences comprised a small fraction of the final dataset (<1% relative abundance on average) and were removed from the study because unlike 16S rRNA amplicons from the 515Y/926R primer sets, analyses of the 18S rRNA amplicons requires overlap between forward and reverse reads for accurate resolution (Yeh et al. 2021), and our sequences were not able to be merged. To ensure accurate representation from the denoised 16S rRNA sequences obtained using only the single-end reads, we evaluated the recovery of taxa and abundances in mock community samples of prokaryotes that were sequenced simultaneously with our environmental samples. Despite the utility and cost-effectiveness of amplicon sequencing, amplicon datasets also present limitations, including biases during PCR which can miss certain groups, especially dinoflagellates (Koumandou et al. 2004), uneven copy numbers of the 16S or 18S rRNA genes across taxa (Decelle et al. 2015), relatively shallow taxonomic resolution (Jovel et al. 2016), and the compositional nature of amplicon sequencing data which requires unique statistical approaches (Gloor et al. 2017). In the context of amplicon sequence data, “phytoplankton” herein refers to ASVs classified as cyanobacteria and eukaryotic plastid sequences, although we recognize that mixotrophic and phagotrophic lifestyles may be included in this broad definition.

Statistical analyses were conducted using the R packages phyloseq v 1.36.0 (McMurdie and Holmes 2013), ggplot2 v 3.4.3 (Wickham 2016), pheatmap v 1.0.12 (Kolde 2019), and microbiome v 1.14.0 (Lahti and Shetty 2017). An Aitchison distance (Aitchison 1982), the Euclidean distance between centered log-ratio (clr)-transformed compositions, was used on the entire quality-controlled dataset of phytoplankton raw read counts using ‘transform’ in the microbiome (Lahti and Shetty 2017). Ward D2 hierarchical clustering using ‘hclust ()’ in the stats base package of R was applied to this matrix to cluster samples with amplicon data into groups and visualized with dendextend v1.17.1 (Galili 2015). ANCOM-BC v 2.6.0 (Lin and Peddada 2024) was used to assess differential abundance patterns of phytoplankton amplicon data across environmental clusters using linear regression models on a bias-corrected log abundance table with different pseudo-counts. To determine whether the differential abundance patterns examined for a taxon are impacted by zero counts, a sensitivity score for each taxon is determined by calculating the proportion of times the p-value is greater than the specified significance level. Divnet v 0.4.0 (Willis and Martin 2020) was used to estimate differences in alpha diversity and test for significance between spatiotemporal groupings. Pearson’s correlation analyses were conducted in corrplot v 0.92 (Wei and Simko 2021).

Lomb Scargle Periodograms (LSP) in the lomb package v 2.1.0 (Ruf 1999) were used to define seasonality among phytoplankton genera by determining the spectrum of frequencies in a dataset: this approach can account for unevenly sampled time-series data and has been previously applied to microbiome time-series analyses (Auladell et al. 2021). Only genera with annual intervals (peak frequency = 1±0.25, p<0.01) as their most significant periodic trend were considered as having seasonality. A starting frequency of 0.16 was used so as to not include periodic components between two consecutive months. An inverse hyperbolic sine transformation (asinh) was conducted on sequence data prior to LSP. Significance (q-values<0.05) was corrected for multiple-testing using data randomization for LSP analyses using fdrtool v 1.2.17 (Strimmer 2008).

### Metagenomic read recruitment

The finer-scale diversity of cyanobacteria can be difficult to resolve with the use of single gene markers (Rocap et al. 2002) and thus metagenomic read recruitment to genome representatives can increase taxonomic resolution (Delmont and Eren 2018; Lee et al. 2019). To investigate the dominant cyanobacteria within and surrounding Kāneʻohe Bay, we conducted metagenomic read recruitment of 32 metagenomes from KByT and 12 previously published from the open ocean Station ALOHA (PRJNA352737; Mende et al. 2017) to 56 cyanobacterial genomes from *Prochlorococcus* (six minor clades) and the three major lineages of the marine *Synecchococcus*/*Cyanobium* lineage [SC 5.1 (14 minor clades), SC 5.2, and SC 5.3] (**Supporting Information Table S2**). SC 5.2 is the only clade with both *Synechococcus* and *Cyanobium* members (Doré et al. 2020).

A contig database of the 56 cyanobacteria isolate genomes was constructed using anvi’o v 8.0 (Eren et al. 2021) following previously described pipelines (Delmont and Eren 2018).

Briefly, Prodigal v2.6.3 (Hyatt et al. 2010) was used to identify open reading frames (ORFs) from the contigs and an anvi’o database was created using ‘anvi-gen-contigs-db’. Metagenomic reads were first quality filtered using an Illumina-utils library v1.4.1 called ‘iu-filterquality- minoche’ (Eren et al. 2013) that uses quality filtering parameters described previously (Minoche et al. 2011). Quality filtered metagenomic reads were competitively mapped with Bowtie2 v2.3.5 (Langmead and Salzberg 2012) to an anvi’o contig database of cyanobacterial isolate genomes. The ‘anvi-profile’ function stored coverage and detection statistics of each cyanobacterial genomes found in the KByT and Station ALOHA metagenomic samples.

To evaluate the distribution of individual genomes, a “detection” metric, the proportion of the nucleotides in a given sequence that are covered by at least one short read, was used to evaluate if a population was present in a metagenomic sample. A detection value of at least 0.25 was used as a criterion to eliminate false positives, when an isolate genome was falsely found within a sample (Utter et al. 2020). Mean coverage q2q3, which refers to the average depth of coverage excluding nucleotide positions with coverages in the 1^st^ and 4^th^ quartiles, was mapped for each genome. Mean coverage q2q3 was summed across all cyanobacterial genomes per sample and then the genome (or all genomes in a clade) was divided by this sum to determine a relative abundance of a genome (or a clade) in each sample. Average nucleotide identity (ANI) was calculated using pyANI v0.2.12 (Pritchard et al. 2015). A phylogenomic tree was estimated from the 56 cyanobacterial isolates and an outgroup using GTotree v1.4.16 (Lee 2019) with cyanobacterial single-copy genes and visualized in FigTree v 1.4.4 (http://tree.bio.ed.ac.uk/).

## Data & code availability

Sequencing data are available in the National Center for Biotechnology Information (NCBI) Sequence Read Archive (SRA) under BioProject number PRJNA706753 as well as PRJNA971314. Environmental data were submitted to BCO-DMO under https://www.bco-dmo.org/project/663665. Code used in the analysis is available at https://github.com/tucker4/Tucker_Phytoplankton_KByT_HeNERR.

## Results

### Biogeochemical parameters

Along the nearshore to open ocean waters of the tropical Pacific, biogeochemical parameters sharply declined across both small spatial scales and vast stretches of ocean (**Supporting Information Table S1)**. On average across time, fluorometrically determined Chla concentrations at stations in the coastal waters of Kāneʻohe Bay increased 18-fold (1.6±1.9 *vs.* 0.09±0.03 µg L^-1^, mean±sd) from the open ocean and 3-fold (1.6±1.9 *vs.* 0.6±0.4 µg L^-1^) from the immediately adjacent offshore waters. The estuarine waters of Heʻeia Fishpond harbored higher Chla concentrations compared to coastal stations (3.9±3.8 *vs.* 1.6±1.9 µg L^-1^, **Fig. 1c**). Mean Chla concentrations increased 43-fold between the estuarine waters of Heʻeia Fishpond and the open ocean (3.9±3.8 *vs.* 0.09±0.03 µg L^-1^, **Supporting Information Table S1**) and 7- fold over the <6 km distance covering the interior and surrounding waters of Kāneʻohe Bay (3.9±3.8 *vs.* 0.6±0.4 µg L^-1^; **Fig. 1c**, **Supporting Information Table S1**). Stations offshore from Kāneʻohe Bay had elevated concentrations of Chla compared to the open ocean (7-fold increase; 0.6±0.4 *vs.* 0.09±0.03 µg L^-1^). Elevated phytoplankton biomass was a persistent feature within Kāneʻohe Bay, with higher Chla concentrations detected in at least one of the stations positioned in the coastal environment compared to the stations offshore during all 36 sampling events (**Supporting Information Fig. S1**).

Elevated concentrations of inorganic nutrients (silicate, nitrate+nitrite, phosphate, ammonia) were also found in the nearshore waters of Kāneʻohe Bay compared to offshore and open ocean stations (**Fig. 1d**, **Supporting Information Table S1**). Mean silicate concentrations at Wai2 of the estuarine stations were 107.33±45.38 µM, compared to 8.79±5.58 µM in the coastal stations (**Fig. 1d**). Across all KByT stations, phosphate and silicate were positively correlated with increasing chlorophyll *a* concentrations (p<0.001, p=0.001, respectively), while nitrate+nitrite concentrations did not correlate with chlorophyll *a* concentrations (p=0.08, **Supporting Information Fig. S2**). Importantly, both inorganic nutrients and chlorophyll *a* concentrations increased with decreasing salinity (**Supporting Information Fig. S2**), suggesting that freshwater and brackish water input (e.g., surface stream and submarine groundwater discharge) may be an important driver of nutrient delivery and subsequent phytoplankton enhancement within Heʻeia Fishpond and Kāneʻohe Bay (McKenzie et al. 2019). Despite the overall increase of inorganic nutrients in the estuarine and coastal stations, all stations were below 16:1 N:P ratios using dissolved inorganic nitrogen (nitrate+nitrite plus ammonia) and phosphate (**Fig. 1e**, (Redfield 1960).

### Microbial cell counts and phytoplankton pigments

In contrast to coastal stations where *Synechococcus* cellular abundance was high, *Prochlorococcus* cellular abundance was elevated in the stations positioned in the offshore waters surrounding Kāneʻohe Bay and at the open ocean Station ALOHA (**Fig. 2a, Supporting Information Table S1**). Cellular abundances of heterotrophic bacteria and photosynthetic picoeukaryotes were also greater in the coastal stations compared to offshore and open ocean (**Fig. 2a, Supporting Information Table S1**). In coastal Kāneʻohe Bay, ratios of fucoxanthin, peridinin, and alloxanthin to total chlorophyll *a* (Tchla) concentrations were higher than in the offshore and in the open ocean, indicating an increase in diatoms, dinoflagellates, and cryptophytes (respectively) closer to shore (**Fig. 2b, Supporting Information Table S3**). In contrast, pigments relative to Tchla for photosynthetic pigments diagnostic of prymnesiophytes (19′-hexanoyloxyfucoxanthin), pelagophytes (19′-butanoyloxyfucoxanthin), cyanobacteria (zeaxanthin), and *Prochlorococcus* (divinyl chlorophyll *a*) were higher in the offshore and in the open ocean compared to the coastal environment (**Fig. 2b**, **Supporting Information Table S3**).

**Fig. 2.**
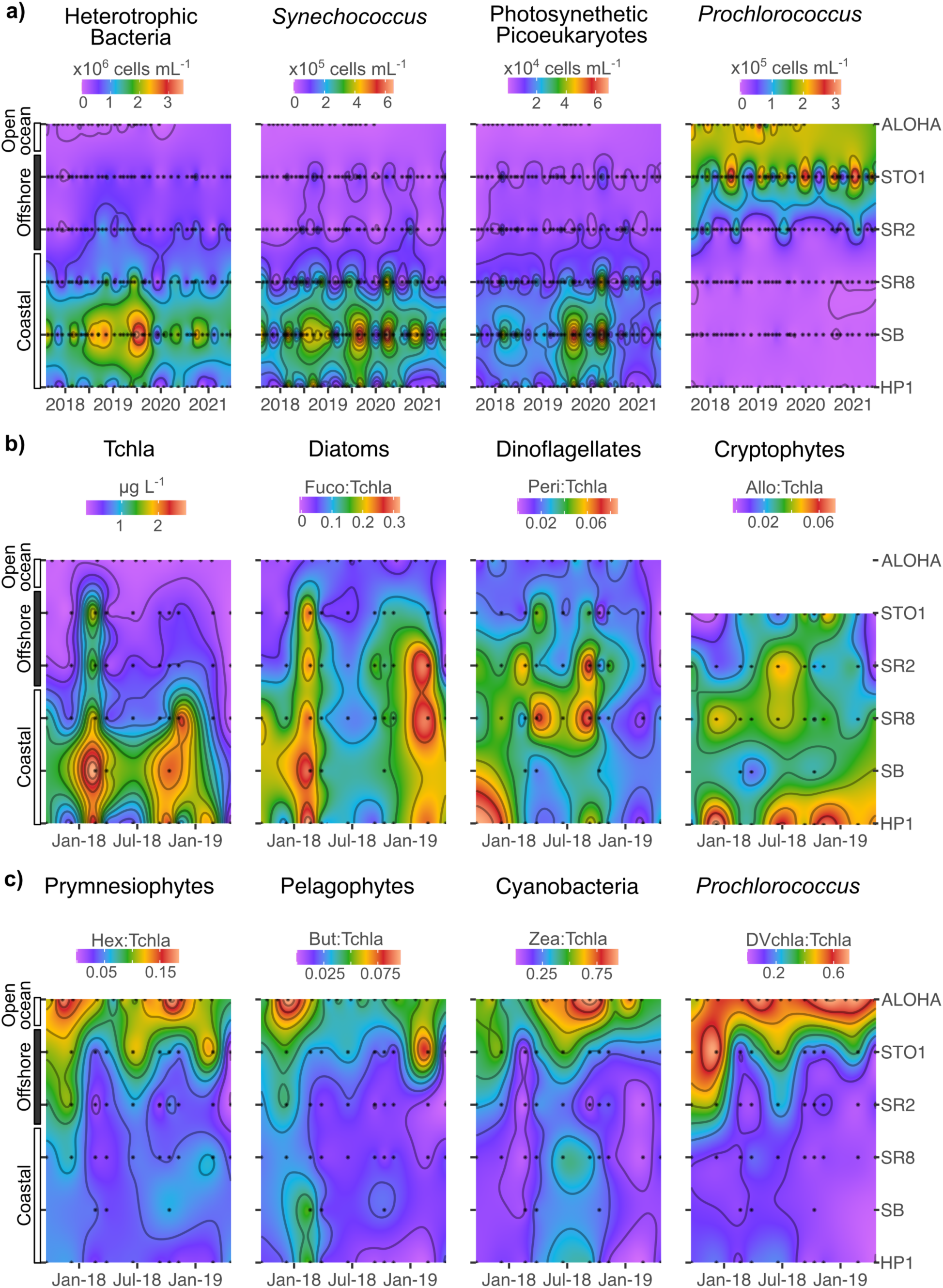
Microbial cellular abundances and pigment concentration vary through time and space across stations from the southern sector of Kāneʻohe Bay (HP1, SB, SR8), offshore stations (SR2 and STO1), and open ocean Station ALOHA: a) Cellular abundances (cells mL^-1^) of heterotrophic bacteria*, Synechococcus*, photosynthetic picoueukaryotes, and *Prochlorococcus*; b) Total chlorophyll *a* (Tchla) concentrations (µg L^-1^) and ratios of phytoplankton pigments indicative of specific phytoplankton groups relative to Tchla. Note, alloxanthin was below detection levels for Station ALOHA and not presented here. Abbreviations: Fuco: Fucoxanthin, Peri: Peridinin; Allo: Alloxanthin; Hex: 19′-hexanoyloxyfucoxanthin; But: 19′-butanoyloxyfucoxanthin; Zea: zeaxanthin; Dvchla: divinyl chlorophyll *a*.

At stations within Kāneʻohe Bay, peaks in the cellular abundances of heterotrophic bacteria, *Synechococcus*, and photosynthetic picoeukaryotes coincide with summer sampling months (**Fig 2b, Supporting Information Fig. S3**). In addition, increased rainfall and wind speeds from storms during three winter sampling events (February 21, 2018, March 3, 2020, and February 12, 2021) aligned with large changes in photosynthetic picoeukaryotes and *Synechococcus* cellular abundances and Chla concentrations (**Fig 2b, Supporting Information Fig. S3**). Elevated Tchla and fucoxanthin (a pigment indicative of diatoms) were also measured during the February 2018 storm in both coastal Kāneʻohe Bay and the adjacent offshore (**Fig 2b**), suggesting that episodic weather events can trigger phytoplankton responses at relatively large spatial scales.

### Phytoplankton community composition through 16S rRNA gene sequencing

We delineated 505 phytoplankton ASVs across 366 samples, including 66 from cyanobacteria and 439 from eukaryotic plastids. Examining the distribution of phytoplankton ASVs across samples revealed that phytoplankton communities clustered into three major community types that coincide with spatial differences in biogeochemistry (**Fig. 3a-b**, **Supporting Information Table S4**). We labeled the three major community types as nearshore, transition, and offshore because of their distinct biogeochemical characteristics and geographic location (**Supporting Information Table S4, Fig. S4**). The nearshore cluster of 229 samples included all samples collected from six stations found most closely located to land (Wai2, Kahoʻokele, SB, CB, HP1, AR) and at least one sample from the six remaining stations. The transition cluster of 85 samples consisted of samples collected from stations not immediately next to land (NB, SR8, NR2, SR2, STO1, NTO1), while the offshore cluster encompassed 52 samples collected exclusively from the four stations located the furthest distance from land (SR2, NR2, STO1, NTO1; **Fig. 3b**). A small number of samples (4 of 72) collected from the two furthest offshore stations (STO1 & NTO1) resolved as a nearshore community type, suggesting that nearshore phytoplankton communities are rarely, but occasionally found outside of the bay (**Fig. 3b**). However, despite a high-level of previously reported water exchange (Lowe et al. 2009), we do not observe offshore phytoplankton communities establishing at stations within Kāneʻohe Bay.

**Figure 3.**
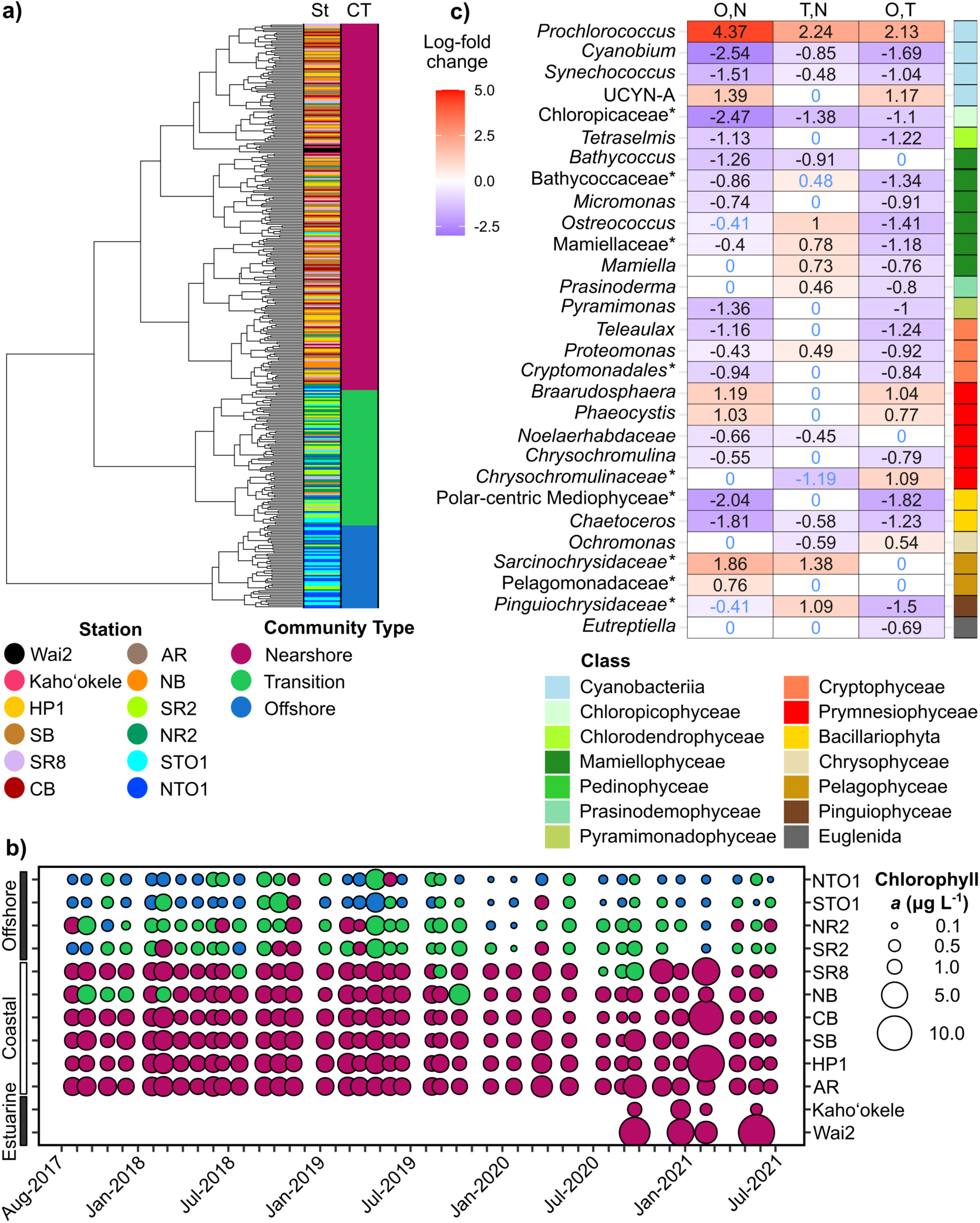
a) Hierarchical clustering of phytoplankton communities from 366 samples collected at 12 sampling stations form three major groups, hereafter referred to as community types: nearshore, transition, and offshore. b) The distribution of samples defined as nearshore, transition, and offshore community types across stations and sampling events (36 total between 2017-2021) shows spatial and temporal persistence in the distinct community types. The size of the circle represents the fluorometric chlorophyll *a* concentration (Chla) during the time of sampling and the color of the circle represents the community type: nearshore, transition, and offshore. c) Log-fold changes in phytoplankton genera with significantly different abundances between the offshore compared to the nearshore (O, N), transition compared to the nearshore (T, N), and the offshore compared to the transition (O, T). Log-fold changes in abundance are shown in text, with black text indicating the comparison passed a sensitivity analysis to assess the impact of pseudo-counts. Phytoplankton groups that were classified at the family-level but unidentified at the genus-level are denoted with an asterisk.

ASV richness was the highest in the nearshore, while the transition community group had the highest Shannon diversity estimate (**Supporting Information Table S5**). A large proportion of the ASVs were shared among all three community types (n=172 of 505), while 126 ASVs were uniquely found in the nearshore, 29 ASVs were found only in the offshore, and only nine ASVs were found solely within the transition community type (**Supporting Information Fig S5**). Differential abundance analysis of amplicon data (**Fig. 3c**) were consistent with results from phytoplankton pigments (**Fig. 2**): the offshore phytoplankton community was enriched in *Prochlorococcus*, prymesiophytes (e.g. *Phaeocystis* and *Braarudosphaera*), and pelagophytes (e.g. *Pelagomonadaceae*, *Sacrinochrysidaceae*) relative to the nearshore, while the nearshore was enriched in diatoms (e.g. *Chaetoceros*, Polar-centric Mediophyceae) and cryptophytes (e.g. *Teleaulax*, *Cryptomonadales*) relative to the offshore (**Fig. 3c; Supporting Information Table S6**). The transition community was uniquely enriched in green algae (e.g. *Ostreococcus*, Mamiellophyceae) and Pinguiophyceae relative to the nearshore and the offshore. However, the transition also represented characteristics of both the nearshore (i.e. similar abundances of diatoms as the nearshore) and offshore communities (i.e. elevated abundances of *Prochlorococcus* compared to the nearshore; **Fig. 3c**).

### Cyanobacterial population structure

Metagenomic read recruitment to 56 genomes of the cyanobacterial genera *Prochlorococcus*, *Synechococcus*, and *Cyanobium* showed that *Prochlorococcus* HLI, HLII, and LLI, and *Synechococcus* SC 5.1 II, WPC2, III, UC-A, and IX and SC 5.3 were detected in metagenomes from the Kāneʻohe Bay Time-series and surface ocean samples from Station ALOHA (**Supporting Information Fig. S6**). Although *Cyanobium* 16S rRNA gene ASVs were detected in the amplicon data, no *Cyanobium* representatives (SC 5.2) were detected in our metagenomic read recruitment. Genomes from *Synechococcus* SC 5.1 II and SC 5.3 and *Prochlorococcus* HLII were among the most abundant representatives within our samples (**Fig. 4**). *Prochlorococcus* HLII comprised 98.9±1.0% (mean±sd) of cyanobacterial relative abundance in the open ocean and 83.1±17.4% of the cyanobacterial relative abundance in the offshore.

**Fig. 4.**
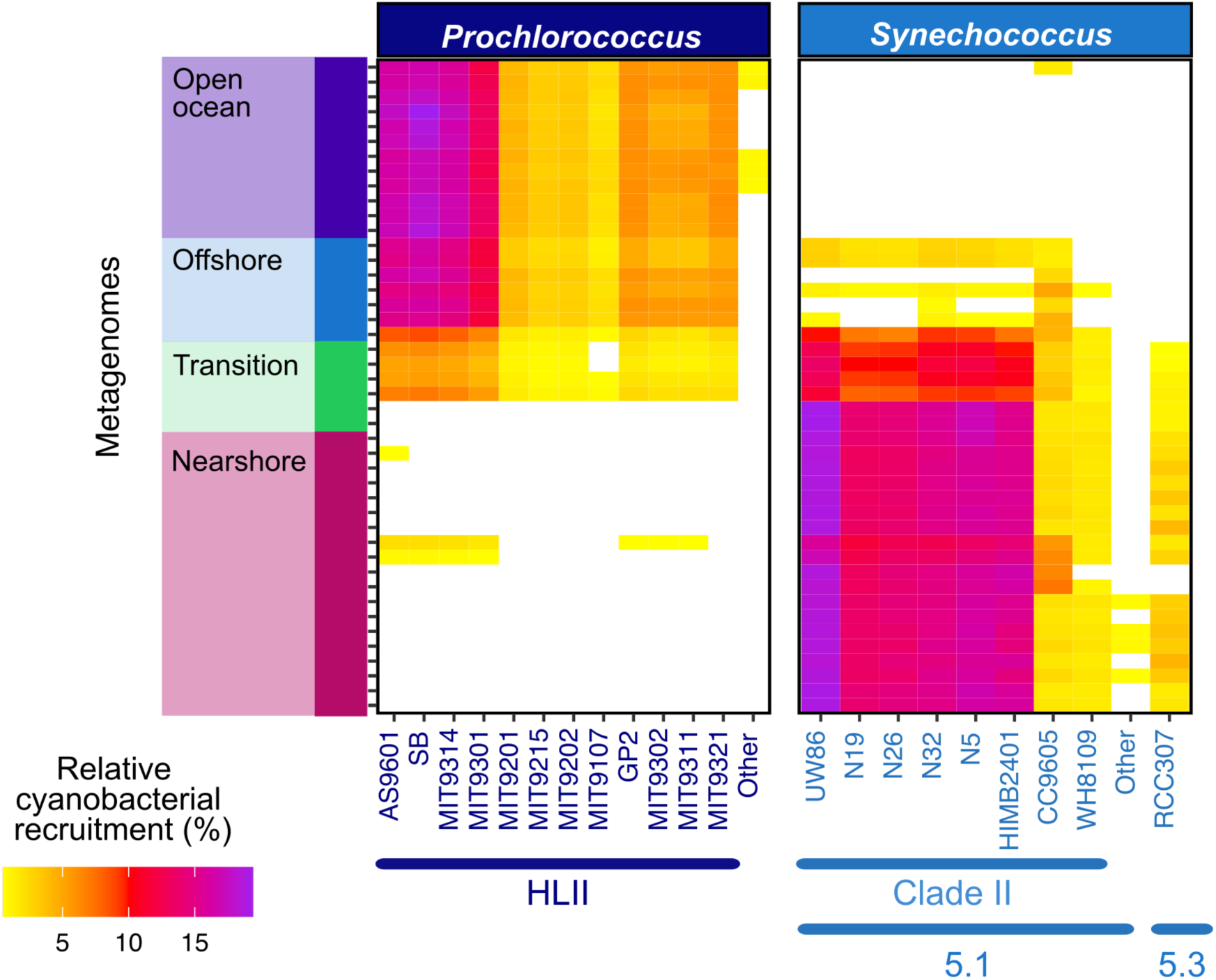
The relative abundance of cyanobacterial genomes across KByT and Station ALOHA metagenomes. *Prochlorococcus* HLII had the highest abundance relative to the other cyanobacteria in the offshore and Station ALOHA metagenomes, while *Synechococcus* clade II (SC 5.1) dominated in the nearshore metagenomes. The other group within *Prochloroccocus* encompassed isolates with low relative abundance from the LLI and HLI clades. The other within *Synechococcus* 5.1 encompassed isolates with low relative abundance from WPC2, III, UC-A, and IX clades. Recruitment of <0.5% not shown.

*Prochlorococcus* HLII recruited only a small proportion of reads from a handful of nearshore samples, where it made up 1.1±2.4% of the cyanobacterial relative abundance. *Synechococcus* clade II comprised 96.3±2.5% of the cyanobacterial relative abundance in the nearshore Kāneʻohe Bay community type. *Synechococcus* SC 5.3 also recruited some metagenomic reads, but only from coastal Kāneʻohe Bay samples and at low relative cyanobacterial abundance (2.4±1.2%) (**Fig. 4**).

Within the *Prochlorococcus* HLII and *Synechococcus* II clades, read recruitment varied between closely related genomes. Read recruitment was substantially higher in *Synechococcus* clade II isolate UW86 compared to all other clade II genomes, despite sharing >95% ANI with most other clade II genomes (**Supporting Information Fig. S6**). Within *Prochlorococcus* HLII, isolate genomes AS9601, SB, MIT9314, and to a lesser extent MIT9301, recruited a substantially greater proportion of reads than other members of this clade. AS9601, SB, MIT9314, and MIT9301 shared 94% ANI, which is higher than what was shared with other HLII isolates (<93%), with the exclusion of MIT9215 and MIT9202 who share 97% ANI with each other (**Supporting Information Fig. S6**).

### Seasonality in biogeochemistry and community composition

We detected seasonal differences in biogeochemistry and phytoplankton community composition across nearshore Kāneʻohe Bay and the adjacent transition and offshore waters, although seasonality in the transition and offshore waters was more subtle than patterns in the nearshore. Surface seawater temperatures were significantly cooler in the winter than in the summer in nearshore Kāneʻohe Bay and the adjacent transition and offshore waters (**Supporting Information Table S7**). Inorganic nutrient concentrations (e.g. phosphate, nitrate+nitrite, silicate) did not significantly change in the offshore between summer and winter months (**Supporting Information Table S7**), however both phosphate and silicate concentrations increased in the summer compared to the winter in the nearshore (**Supporting Information Table S7**). Cellular concentrations of *Synechococcus* and heterotrophic bacteria were higher in the summer than the winter in the nearshore, transition, and offshore (**Fig. 5a**, **Supporting Information Table S7**). Photosynthetic picoeukaryote cellular concentrations also increased during the summer in coastal and transition environments (coastal: p=0.004, transition: p<0.001, **Fig. 5a, Supporting Information Table S7**), but did not vary seasonally in the offshore. In contrast, *Prochlorococcus* cell concentrations only varied seasonally in the nearshore where it increased in the winter (p<0.001, **Supporting Information Table S7**).

**Fig. 5.**
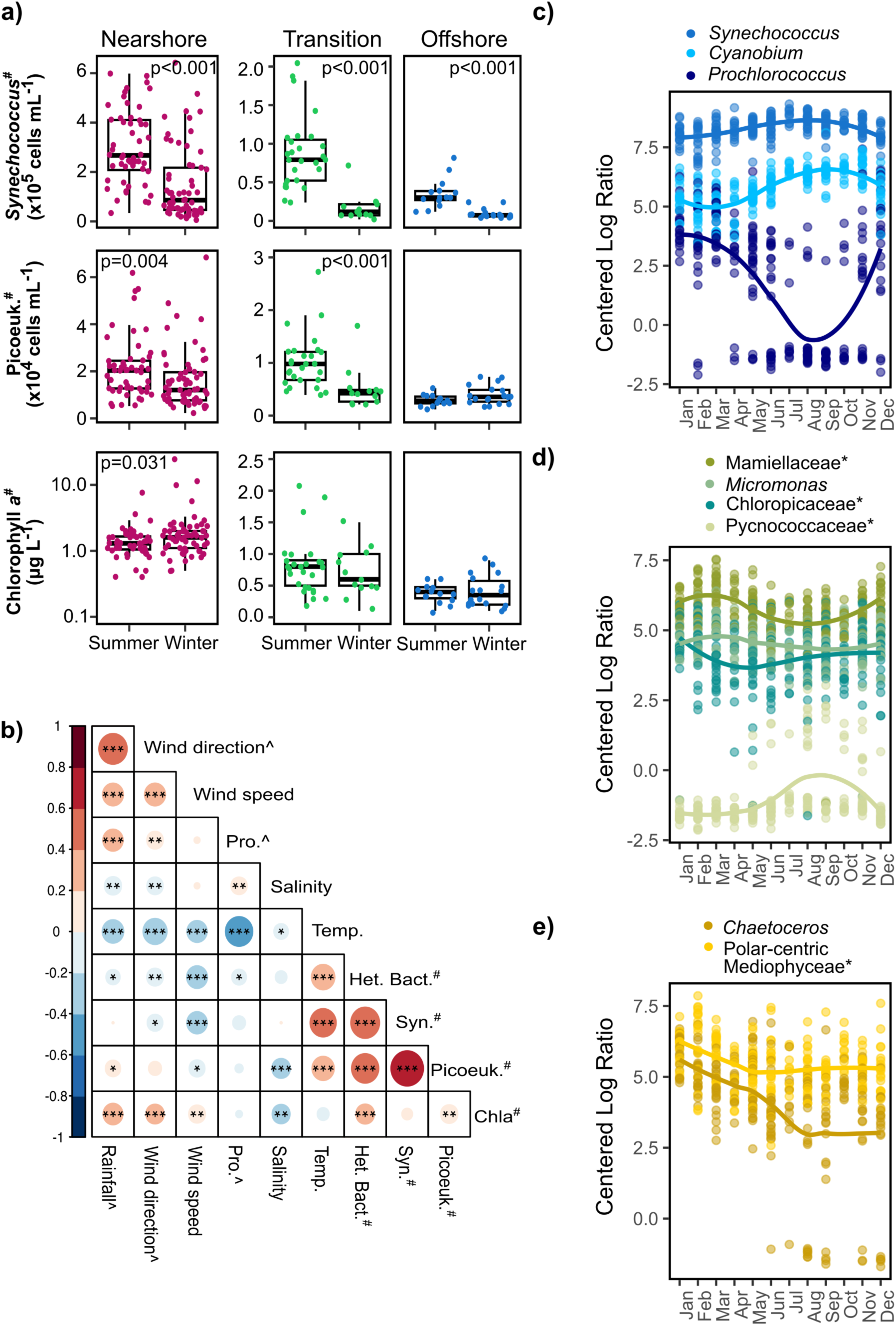
a) Changes in cell counts of *Synechococcus* and photosynthetic picoeukaryotes and fluorometric chlorophyll *a* concentrations (Chla) across seasons and environments. Note, that Chla concentrations are presented on a log scale for the nearshore, but not for the transition and offshore environments. b) Correlations between Chla and other environmental parameters in the nearshore. c-e) Genera belonging to cyanobacteria (c), archaeplastida (green algae) (d), and stramenopiles (diatoms) (e) show seasonal changes in abundance in the nearshore. A local polynomial regression fit line is shown for each genus. Phytoplankton groups that were classified at the family-level but unidentified at the genus-level are denoted with an asterisk. A pound sign (#) denotes variables with log transformations, while a carrot (^) denotes variables with log+1 transformations. Het.Bac: heterotrophic bacteria (cells mL^-1^); Syn: *Synechococcus* (cells mL^-1^); Pro: *Prochlorococcus* (cells mL^-1^); Picoeuk: Photosynthetic picoeukaryotes (cells mL^-1^); Temp: Seawater temperature (°C); Chla: Chlorophyll *a* (µg L^-1^); Salinity (ppt); Wind direction (degrees); Wind Speed (ms^-1^); Rainfall (mm).

Chla concentrations increased in the winter compared to the summer in the nearshore (mean±sd; winter: 2.3±3.4 µg L^-1^, summer: 0.8±0.5 µg L^-1^; p=0.031; **Fig. 5a, Supporting Information Table S7**), but not in the transition or offshore waters. Importantly, given that the collection method for Chla concentrations likely missed most small cyanobacteria, it is possible that seasonality has been underestimated. Nearshore Chla concentrations increased with wind speed, wind direction, and rainfall, but not with seawater temperature (**Fig. 5b**). Ratios of phytoplankton pigments to Tchla representing diatoms and pelagophytes showed a negative correlation with seawater temperature in the nearshore community type, suggestive that these phytoplankton may be driving the winter increase in Chla concentrations in the nearshore (**Supporting Information Fig. S7**). In contrast, pigments indicative of cyanobacteria showed significant positive relationship with seawater temperature in the nearshore (**Supporting Information Fig. S7**). Correlations between seawater temperature and pigment to Tchla ratios were not detected in the transition and offshore cluster (**Supporting Information Fig. S7**). Chla concentrations in the transition and offshore also did not correlate with seawater temperatures, but did positively correlate with rainfall (**Supporting Information Fig. S8**).

Multi-year seasonality was observed among 23 phytoplankton genera (or groups unclassified at the genus-level, but classified at the family-level) within each of the clusters, including 18 nearshore, 5 in the transition, and 6 offshore (Lomb Scargle Periodogram test, q ≤ 0.5, **Supporting Information Table S8**). *Synechococcus*, *Cyanobium*, *Crocosphaera*, and unidentified Pycnococcaceae increased in abundance in the summer months (**Supporting Information Fig. S9**). However, most genera with seasonal patterns peaked in abundance during the winter, including almost all belonging to diatoms (e.g. *Chaetoceros* and unclassified polar-centric Mediophyceae), green algae (e.g. unclassified Mamiellophyceae, *Micromonas*, and *Mamiella*), prymnesiophytes (e.g. *Isochrysis*, *Chrysochromulina*, and *Braarudosphaera*), and Dictyochophyceae (*Helicopedinella*, *Mesopedinella*) (**Supporting Information Fig. S9**). ASV richness and Shannon diversity increased in the winter compared to the summer in all three community types as well (**Supporting Information Table S9**).

In the nearshore, dominant phytoplankton groups with seasonal patterns include cyanobacteria, green algae, and diatoms (**Fig. 5 c, d, e**). While the two diatom groups are both enriched in the winter months relative to the summer months, the seasonality of members belonging to green algae and cyanobacteria differs by genus in the nearshore (**Fig. 5 c, d, e**). The distinct seasonal patterns among the cyanobacteria, such that *Prochlorococcus* increases in the winter in the nearshore, while *Synechococcus* increases in the summer, is further supported by cellular abundance data (**Supporting Information Table S7**). As water sample collection included a pre-filtering step with 85 μm mesh to exclude zooplankton, our estimates may have resulted in an underestimation of diatom species larger than 85 μm.

## Discussion

The activities and distributions of phytoplankton have important implications for food web dynamics, biocultural restoration, and adaptive management of near-island waters of the tropical Pacific, especially under predicted climate change conditions that are expected to shift nutrient availability, phytoplankton size, productivity, and biomass in open oceans (Boyce et al. 2010; Flombaum et al. 2020; Li et al. 2020). Ecosystem-level time-series analyses spanning estuarine waters within Kāneʻohe Bay of the Hawaiian island of Oʻahu to the North Pacific Subtropical Gyre revealed that surface ocean biogeochemistry, phytoplankton biomass, and phytoplankton community structure varied dramatically across both broad (e.g. nearshore Kāneʻohe Bay to the NPSG) and narrow (e.g. nearshore Kāneʻohe Bay to adjacent offshore waters) spatial scales. Through investigations of these spatiotemporal dynamics, we gained insight into the ecology and phenology of phytoplankton communities of surface oceans in the tropical Pacific and identified indicators to help evaluate deviations from current conditions.

### Distinct phytoplankton and biogeochemical regimes within and adjacent to Kāneʻohe Bay

In line with previous work, our results show that the elevated phytoplankton biomass in nearshore Kāneʻohe Bay is predominantly driven by the freshwater input of land-derived and anthropogenic nutrients delivered via surface streams to coastal waters (Cox et al. 2006; Drupp et al. 2011; Yeo et al. 2013) and submarine groundwater-derived inputs (Dulai et al. 2016; Ghazal et al. 2023), which reduces the prevailing nitrogen limitation and enhances phytoplankton growth (Ringuet and Mackenzie 2005; Carlo et al. 2007). Although not examined in this study, sewage effluent outfall (Laws and Redalje 1982; Laws and Taguchi 2018), ocean-driven internal waves (Gove et al. 2016; Comfort et al. 2024), and coral-reef processes (James et al. 2020) are other likely important mechanisms for nutrient delivery into coastal Kāneʻohe Bay. Over the four year time-series, inorganic nutrients and phytoplankton biomass were persistently elevated within Kāneʻohe Bay relative to offshore stations, with enhanced phytoplankton biomass primarily attributed to *Synechococcus* and diatoms. Members of phytoplankton groups with larger cell sizes were enriched in the nearshore relative to the offshore, including diatoms, dinoflagellates, and cryptophytes, following previous work that has observed increased phytoplankton cell sizes with increased nutrient availability (Chisholm 1992; Hillebrand et al. 2022). Offshore phytoplankton communities were enriched in relatively smaller phytoplankton groups, including *Prochlorococcus*, prymnesiophytes and pelagophytes, which have been reported to reach high abundances in warm oligotrophic oceans (Cuvelier et al. 2010; Li et al. 2013; Rii et al. 2016).

Despite a constant input of oceanic seawater, phytoplankton communities within Kāneʻohe Bay were structured into distinct communities that varied in composition, diversity, and potentially lifestyles. Limited dispersal alone cannot explain this strong structuring, as there is a high overlap in ASVs shared across the community types and most of our sampling stations in the Bay receive advection of oceanic-sourced waters (Lowe et al. 2009). Instead, community structure most likely reflects a high degree of environmental selection for locally adapted populations across this sharp environmental gradient (Yeo et al., 2013; Tucker et al., 2021). One potential outcome of environmental selection and resource competition is the enrichment of phytoplankton associated with mixotrophic lifestyles closer to shore, including *Teleaulax* (Cryptophyta), *Pyamimonas* (Chlorophyta), *Tetraselmis* (Chlorophyta), and dinoflagellates (Stoecker et al. 2016). By supplementing primary production with heterotrophy, mixotrophy serves as a source of inorganic and organic nutrients, increases carbon transfer to high trophic levels, and shifts the community toward larger plankton size classes (Stoecker et al. 2016; Ward and Follows 2016). While establishing the relative importance of mixotrophy across nearshore Kāneʻohe Bay to the adjacent offshore environment requires further investigation, the abundant bacterial prey items and limited inorganic nutrients of coastal Kāneʻohe Bay may favor mixotrophy over autotrophy (Edwards 2019). These initial insights highlight important implications for the structuring of phytoplankton communities across the nearshore to open ocean waters of the tropical Pacific, including distinctions in food web dynamics, phytoplankton ecologies, and contributors to primary productivity.

### Drivers of seasonality in nearshore chlorophyll a concentrations

In Hawaiʻi, storm events increase during the winter months leading to higher stream discharge, increased nutrient input, and phytoplankton growth (Ringuet and Mackenzie 2005; Cox et al. 2006; Yeo et al. 2013). We observed three distinct storm events within winter months that likely contributed to the observed increase in phytoplankton biomass during wet (winter) months in the nearshore. Previous work has shown that the wet (winter) season has strong impacts on the food web dynamics of Kāneʻohe Bay: increased diatom biomass decreases the number of trophic levels between autotrophs and metazoan larvae and nauplii and leads to an increase in total community biomass and in the transfer of production to metazoans (Selph et al. 2018). Periods of high nutrient availability favors diatoms because they are fast-growing nutrient opportunists with high maximum growth but low affinity for uptake (Laws 1975; Dell’Aquila et al. 2017; Inomura et al. 2023). In contrast, in the summer under high-light, warm-water temperatures and limited nutrients, the high affinity for uptake but low maximum growth of *Synechococcus* is expected to be favored (Agawin et al. 1998, 2000; Domínguez-Martín et al. 2022). These trade-offs between growth and nutrient uptake strategies, along with changes in the environment that favors one strategy over another (Edwards et al. 2012, 2024), likely produced the dynamic seasonal patterns in the dominant phytoplankton observed in nearshore Kāneʻohe Bay.

#### Phytoplankton indicators of climate change impacts

Biodiversity within phytoplankton groups can modulate the impacts of variable temperature and nutrient regimes on growth, and thus may provide good bioindicators of resilience to ongoing climate impacts (Martiny et al. 2022; Vrana et al. 2023). Within *Prochlorococcus* and *Synechococcus*, small gene gain and loss events have allowed for local adaptations of major and minor clades to variable nutrient concentrations and substrates (Berube et al. 2019; Yan et al. 2022; Doré et al. 2023; Ustick et al. 2023). We find the dominance of *Synechococcus* clade II and *Prochlorococcus* HLII in coastal and offshore environments of the tropical Pacific, respectively, as well as clear population-level differences within these two clades. Using read recruitment from global surface and subsurface ocean metagenomes, Delmont & Eren (2018) found distinct distribution patterns among similar subgroups of HLII isolates as those reported here and associated such differences in fitness with fine-scale differences in gene content. *Prochlorococcus* HLII from the offshore waters adjacent to Kāneʻohe Bay and Station ALOHA showed high similarity in population structure and thus likely represent continuous populations with ongoing gene flow and adaptations driven by similar environmental pressures. Continued examination of population structure, as well as cellular and relative abundances of these cyanobacteria, could identify eventual expansions of ultraoligotrophic waters into Kāneʻohe Bay, shifts in gene-flow, gains or losses of genes related to nutrient acquisition or temperature stress, and selection for clades with unique ecological adaptations.

In addition, deviations in the magnitude and timing of seasonal changes in *Synechococcus* and diatom abundances could also be used to identify variable responses to warming temperatures and changing nutrient availability. Temperature has been shown to positively impact *Synechococcus* division rates (Hunter-Cevera et al. 2016), suggesting that under warmer seawater temperatures *Synechococcus* may further dominate the nearshore phytoplankton community. In contrast, while some diatom species appear to have the capacity to adapt physiologically and genetically to increased temperatures (Schaum et al. 2017; Agusti et al. 2019; Hattich et al. 2024), temperature adaptation may be limited under the low nitrogen concentrations that are characteristic of Kāneʻohe Bay (Li et al. 2018; Aranguren-Gassis et al. 2019). The magnitude and frequency of storm events on windward Oʻahu are expected to change with ongoing climate change (Zhang et al. 2016; Fandrich et al. 2022), suggesting that it is possible that under wetter conditions nearshore diatoms may keep pace with increasing temperatures, but under drier conditions increased nitrogen-limitation may lead to reduced diversity, seasonality, and/or overall abundances of diatoms. It has also been observed that high-island environments of Hawaiian Islands promote elevated delivery of submarine groundwater discharge into nearshore regions during king tides (McKenzie et al. 2021), thus further impacting phytoplankton community composition under combined scenarios of increased storms, stratification, and sea-level rise.

#### Implications for biocultural restoration and adaptive management

Within Heʻeia and across the islands of Hawaiʻi, Indigenous People and Local Communities are undertaking biocultural restoration projects of Indigenous aquaculture systems to maximize primary and secondary production of the estuarine environment, including removing invasive mangrove, managing stream use to ensure adequate flow and water quality, and engineering water exchange through repairs and updates to fishpond walls (Möhlenkamp et al. 2019). The baseline understanding of phytoplankton communities provided by this study helps to inform biocultural stewards of Heʻeia and Kāneʻohe Bay of the conditions that promote phytoplankton growth and subsequently herbivorous fish growth. Despite differences in biogeochemistry across nearshore stations, phytoplankton communities sampled from the Heʻeia Fishpond and across the bay grouped as one nearshore community type. This emphasizes the connectedness of these estuarine and coastal systems and highlights the need to manage them in close coordination. Further growing this baseline understanding of phytoplankton assemblages to relate to the timing and conditions documented in Hawaiian knowledge systems like *kaulana mahina*, the Hawaiian lunar calendar (Nuʻuhiwa 2019), and in regard to life cycles of bioculturally relevant species like ʻamaʻama (*Mugil cephalus*) which are grown in the fishpond and feed on diatoms and other microphytoplankton at juvenile stages (Hiatt 1947), would advance this area of study and utility for management within these Indigenous aquaculture systems. Our study also provides opportunities to refine hydrological models conducted in social-ecological systems (Ghazal et al. 2023) to further inform biocultural restoration towards adaptive management.

Despite a poor understanding of the outcomes, near-island food webs will likely shift with ongoing climate change impacts on phytoplankton communities and biomass. Collaborating across diverse knowledge systems to co-produce new knowledge could improve the ability for local people to document and adapt to changes in near-island marine resources (Winter et al. 2020b). Our collaboratively developed study reveals distinct spatial and seasonal dynamics that define phytoplankton communities and biogeochemical conditions from an estuarine aquaculture system on the coast of Oʻahu, Hawaiʻi, to the open ocean of the North Pacific Subtropical Gyre. Understanding the seasonal and spatial dynamics underlying phytoplankton communities and biogeochemistry of near-island environments in the tropical Pacific provides the necessary knowledge to further co-develop capacities to model and track changes to the marine food web, and to build resilience now and in the future.

## Acknowledgements

We thank Paepae o Heʻeia and the Heʻeia National Estuarine Research Reserve for their involvement in and perspectives on the directions on this research. We thank Jason Jones, Ciara Ratum, Hanako Mochimaru, Oscar Ramfelt, Amber Boettiger, Clarisse E.S. Sullivan, Elizabeth A. Monaghan, and Evan Freel for their assistance collecting samples, Gus Robertson for assistance with sonde calibration, Andrew McGowan for help with the creation of maps, and Karen Selph for flow cytometry measurements. Any opinions, findings, and conclusions or recommendations expressed in this material are those of the author(s) and do not necessarily reflect the views of the National Science Foundation or the National Oceanic and Atmospheric Administration (NOAA). This research was funded by the National Oceanic and Atmospheric Administration (NOAA) Margaret A. Davidson Fellowship (No. NA20NOS4200123), National Science Foundation Graduate Research Fellowship Program under Grant No. 1842402, the Colonel Willys E. Lord, DVM and Sadina L. Lord Scholarship, and the University of Hawaiʻi at Mānoa Ecology, Evolution, and Conservation Biology’s Maybelle Roth ARCS Award to SJ Tucker and NSF grants OCE-1538628 and OCE-2149128 to MS Rappé. This is SOEST contribution XXX and HIMB contribution XXX.

## Data availability statement

Data supporting the results within the manuscript are available within the main text (See Materials & Methods). Sequencing data are available in the National Center for Biotechnology Information (NCBI) Sequence Read Archive (SRA) under BioProject number PRJNA706753 as well as PRJNA971314. Environmental data were submitted to BCO-DMO under https://www.bco-dmo.org/project/663665. Code used in the analysis is available at https://github.com/tucker4/Tucker_Phytoplankton_KByT_HeNERR.

## Citations and References

Acevedo-Trejos, E., G. Brandt, J. Bruggeman, and A. Merico. 2015. Mechanisms shaping size structure and functional diversity of phytoplankton communities in the ocean. Sci. Rep. 5: 8918. doi:10.1038/srep08918

Agawin, N., C. Duarte, and S. Agustí. 1998. Growth and abundance of Synechococcus sp. in a Mediterranean Bay: seasonality and relationship with temperature. Mar. Ecol. Prog. Ser. 170: 45–53. doi:10.3354/meps170045

Agawin, N. S. R., C. M. Duarte, and S. Agustí. 2000. Nutrient and temperature control of the contribution of picoplankton to phytoplankton biomass and production. Limnol. Oceanogr. 45: 591–600. doi:10.4319/lo.2000.45.3.0591

Agusti, S., L. M. Lubián, E. Moreno-Ostos, M. Estrada, and C. M. Duarte. 2019. Projected changes in photosynthetic picoplankton in a warmer subtropical ocean. Front. Mar. Sci. 5: 506. doi:10.3389/fmars.2018.00506

Aitchison, J. 1982. The statistical analysis of compositional data. J Royal Statistical Soc Ser B Methodol 44: 139–160. doi:10.1111/j.2517-6161.1982.tb01195.x

Aranguren-Gassis, M., C. T. Kremer, C. A. Klausmeier, and E. Litchman. 2019. Nitrogen limitation inhibits marine diatom adaptation to high temperatures. Ecol. Lett. 22: 1860–1869. doi:10.1111/ele.13378

Auladell, A., A. Barberán, R. Logares, E. Garcés, J. M. Gasol, and I. Ferrera. 2021. Seasonal niche differentiation among closely related marine bacteria. Isme J 1–12. doi:10.1038/s41396-021-01053-2

Azam, F., T. Fenchel, J. Field, J. Gray, L. Meyer-Reil, and F. Thingstad. 1983. The ecological role of water-column microbes in the sea. Mar Ecol Prog Ser 10: 257–263. doi:10.3354/meps010257

Berube, P. M., A. Rasmussen, R. Braakman, R. Stepanauskas, and S. W. Chisholm. 2019. Emergence of trait variability through the lens of nitrogen assimilation in Prochlorococcus. Elife 8. doi:10.7554/elife.41043

Bidigare, R., O. Schofield, and B. Prezelin. 1989. Influence of zeaxanthin on quantum yield of photosynthesis of Synechococcus clone WH7803 (DC2). Mar Ecol Prog Ser 56: 177–188. doi:10.3354/meps056177

Bolyen, E. and others. 2019. Reproducible, interactive, scalable and extensible microbiome data science using QIIME 2. Nat Biotechnol 37: 852–857. doi:10.1038/s41587-019-0209-9

Bopp, L. and others. 2013. Multiple stressors of ocean ecosystems in the 21st century: projections with CMIP5 models. Biogeosciences 10: 6225–6245. doi:10.5194/bg-10-6225-2013

Boyce, D. G., M. R. Lewis, and B. Worm. 2010. Global phytoplankton decline over the past century. Nature 1–6. doi:10.1038/nature09268

Callahan, B. J., J. Wong, C. Heiner, S. Oh, C. M. Theriot, A. S. Gulati, S. K. McGill, and M. K. Dougherty. 2019. High-throughput amplicon sequencing of the full-length 16S rRNA gene with single-nucleotide resolution. **4**: e2492-28. doi:10.1101/392332

Carlo, E. H. D., D. J. Hoover, C. W. Young, R. S. Hoover, and F. T. Mackenzie. 2007. Impact of storm runoff from tropical watersheds on coastal water quality and productivity. Appl. Geochem. 22: 1777–1797. doi:10.1016/j.apgeochem.2007.03.034

Chisholm, S. W. 1992. Phytoplankton size, p. 213–237. In Falkowski, P.G., Woodhead, A.D., Vivirito, K. (eds) Primary Productivity and Biogeochemical Cycles in the Sea. Environmental Science Research, vol 43. Springer, Boston, MA. 10.1007/978-1-4899-0762-2_12

Cloern, J. E. 1996. Phytoplankton bloom dynamics in coastal ecosystems: A review with some general lessons from sustained investigation of San Francisco Bay, California. Rev Geophys 34: 127–168. doi:10.1029/96rg00986

Comfort, C. M., C. Ostrander, C. E. Nelson, D. M. Karl, and M. A. McManus. 2024. A 7-yr spatial time series resolves the island mass effect and associated shifts in picocyanobacteria abundances near O’ahu, Hawai’i. Limnol. Oceanogr. doi:10.1002/lno.12711

Comstock, J., C. E. Nelson, A. James, E. Wear, N. Baetge, K. Remple, A. Juknavorian, and C. A. Carlson. 2022. Bacterioplankton communities reveal horizontal and vertical influence of an Island Mass Effect. Environ Microbiol. doi:10.1111/1462-2920.16092

Cox, E. F., M. Ribes, and I. K. RA. 2006. Temporal and spatial scaling of planktonic responses to nutrient inputs into a subtropical embayment. Mar. Ecol. Prog. Ser. 324: 19–35. doi:10.3354/meps324019

Cuvelier, M. L. and others. 2010. Targeted metagenomics and ecology of globally important uncultured eukaryotic phytoplankton. Proc. Natl. Acad. Sci. 107: 14679–14684. doi:10.1073/pnas.1001665107

Decelle, J. and others. 2015. PhytoREF: a reference database of the plastidial 16S rRNA gene of photosynthetic eukaryotes with curated taxonomy. Mol Ecol Resour 15: 1435–1445. doi:10.1111/1755-0998.12401

Dell’Aquila, G., M. I. Ferrante, M. Gherardi, M. C. Lagomarsino, M. R. d’Alcalà, D. Iudicone, and A. Amato. 2017. Nutrient consumption and chain tuning in diatoms exposed to storm-like turbulence. Sci. Rep. 7: 1828. doi:10.1038/s41598-017-02084-6

Delmont, T. O., and A. M. Eren. 2018. Linking pangenomes and metagenomes: the Prochlorococcus metapangenome. PeerJ 6: e4320. doi:10.7717/peerj.4320

Domínguez-Martín, M. A., A. López-Lozano, Y. Melero-Rubio, G. Gómez-Baena, J. A. Jiménez-Estrada, K. Kukil, J. Diez, and J. M. García-Fernández. 2022. Marine Synechococcus sp. strain WH7803 shows specific adaptative responses to assimilate nanomolar concentrations of nitrate. Microbiol. Spectr. 10: e00187–22. doi:10.1128/spectrum.00187-22

Doré, H. and others. 2020. Evolutionary mechanisms of long-term genome diversification associated with niche partitioning in marine picocyanobacteria. Front Microbiol 11: 567431. doi:10.3389/fmicb.2020.567431

Doré, H. and others. 2023. Differential global distribution of marine picocyanobacteria gene clusters reveals distinct niche-related adaptive strategies. ISME J. 17: 720–732. doi:10.1038/s41396-023-01386-0

Doty, M. S., and M. Oguri. 1956. The island mass effect. J. Cons. perm. int. Explor. Mer 22: 33–37. doi:10.1093/icesjms/22.1.33

Drupp, P., E. H. D. Carlo, F. T. Mackenzie, P. Bienfang, and C. L. Sabine. 2011. Nutrient inputs, phytoplankton response, and CO2 variations in a semi-enclosed subtropical embayment, Kaneohe Bay, Hawaii. Aquat Geochem 17: 473–498. doi:10.1007/s10498-010-9115-y

Dulai, H., A. Kleven, K. Ruttenberg, R. Briggs, and F. Thomas. 2016. Evaluation of submarine groundwater discharge as a coastal nutrient source and its role in coastal groundwater quality and quantity. pp. 187–221. In A. Fares (ed.) Emerging Issues in Groundwater Resources. doi:10.1007/978-3-319-32008-3_8

Dutkiewicz, S., P. Cermeno, O. Jahn, M. J. Follows, A. E. Hickman, D. A. A. Taniguchi, and B. A. Ward. 2020. Dimensions of marine phytoplankton diversity. Biogeosciences 17: 609–634. doi:10.5194/bg-17-609-2020

Edwards, K. F. 2019. Mixotrophy in nanoflagellates across environmental gradients in the ocean. Proc. Natl. Acad. Sci. 116: 6211–6220. doi:10.1073/pnas.1814860116

Edwards, K. F., Y. M. Rii, Q. Li, L. M. Peoples, M. J. Church, and G. F. Steward. 2024. Trophic strategies of picoeukaryotic phytoplankton vary over time and with depth in the North Pacific Subtropical Gyre. Environ. Microbiol. 26: e16689. doi:10.1111/1462-2920.16689

Edwards, K. F., M. K. Thomas, C. A. Klausmeier, and E. Litchman. 2012. Allometric scaling and taxonomic variation in nutrient utilization traits and maximum growth rate of phytoplankton. Limnol. Oceanogr. 57: 554–566. doi:10.4319/lo.2012.57.2.0554

Eren, A. M. and others. 2021. Community-led, integrated, reproducible multi-omics with anvi’o. Nat Microbiol 6: 3–6. doi:10.1038/s41564-020-00834-3

Eren, A. M., J. H. Vineis, H. G. Morrison, and M. L. Sogin. 2013. A filtering method to generate high quality short reads using Illumina paired-end technology. Plos One 8: e66643. doi:10.1371/journal.pone.0066643

Estrada, M. and others. 2016. Phytoplankton across tropical and subtropical regions of the Atlantic, Indian and Pacific Oceans. PLoS ONE 11: e0151699. doi:10.1371/journal.pone.0151699

Falkowski, P. G., R. T. Barber, and V. Smetacek. 1998. Biogeochemical controls and feedbacks on ocean primary production. Science 281: 200–206. doi:10.1126/science.281.5374.200

Fandrich, K. M., O. E. Timm, C. Zhang, and T. W. Giambelluca. 2022. Dynamical downscaling of near-term (2026–2035) climate variability and change for the main Hawaiian Islands. J. Geophys. Res.: Atmos. 127. doi:10.1029/2021jd035684

Finley, A., B. Sudipto, and Hjelle. 2017. MBA: Multilevel B-Spline Approximation.

Flombaum, P., and A. C. Martiny. 2021. Diverse but uncertain responses of picophytoplankton lineages to future climate change. Limnol Oceanogr. doi:10.1002/lno.11951

Flombaum, P., W.-L. Wang, F. W. Primeau, and A. C. Martiny. 2020. Global picophytoplankton niche partitioning predicts overall positive response to ocean warming. Nat. Geosci. 13: 116–120. doi:10.1038/s41561-019-0524-2

Friedrich, T., B. S. Powell, C. A. Stock, L. Hahn-Woernle, R. Dussin, and E. N. Curchitser. 2021. Drivers of phytoplankton blooms in Hawaii: A regional model study. J Geophys Res Oceans 126. doi:10.1029/2020jc017069

Fu, W., J. T. Randerson, and J. K. Moore. 2016. Climate change impacts on net primary production (NPP) and export production (EP) regulated by increasing stratification and phytoplankton community structure in the CMIP5 models. Biogeosciences 13: 5151–5170. doi:10.5194/bg-13-5151-2016

Galili, T. 2015. dendextend: an R package for visualizing, adjusting and comparing trees of hierarchical clustering. Bioinformatics 31: 3718–3720. doi:10.1093/bioinformatics/btv428

Ghazal, K. A., O. T. Leta, and H. Dulai. 2023. Spatiotemporal estimation of fresh submarine groundwater discharge across the coastal shorelines of Oahu Island, Hawaii. Blue-Green Syst. 5: 28–40. doi:10.2166/bgs.2023.010

Gloor, G. B., J. M. Macklaim, V. Pawlowsky-Glahn, and J. J. Egozcue. 2017. Microbiome datasets are compositional: and this is not optional. Front Microbiol **8**: 2224. doi:10.3389/fmicb.2017.02224

Gove, J. M. and others. 2016. Near-island biological hotspots in barren ocean basins. Nature Communications 7: 10581–8. doi:10.1038/ncomms10581

Group, K. N. W. 2021. Kūlana Noiʻi v. 2. University of Hawaiʻi Sea Grant College Program.

Guillou, L. and others. 2013. The protist ribosomal reference database (PR2): a catalog of unicellular eukaryote small sub-unit rRNA sequences with curated taxonomy. Nucleic Acids Res 41: D597–D604. doi:10.1093/nar/gks1160

Haber, M. and others. 2022. Spatiotemporal variation of microbial communities in the ultra-oligotrophic eastern Mediterranean Sea. Front Microbiol 13: 867694. doi:10.3389/fmicb.2022.867694

Hagstrom, G. I., C. A. Stock, J. Y. Luo, and S. A. Levin. 2024. Impact of dynamic phytoplankton stoichiometry on global scale patterns of nutrient limitation, nitrogen fixation, and carbon export. Glob. Biogeochem. Cycles 38. doi:10.1029/2023gb007991

Hattich, G. S. I. and others. 2024. Temperature optima of a natural diatom population increases as global warming proceeds. Nat. Clim. Chang. 14: 518–525. doi:10.1038/s41558-024-01981-9

Henson, S. A., B. B. Cael, S. R. Allen, and S. Dutkiewicz. 2021. Future phytoplankton diversity in a changing climate. Nat Commun 12: 5372. doi:10.1038/s41467-021-25699-w

Hiatt, R. 1947. Food-chains and the food cycle in Hawaiian fish ponds.–part I. The food and feeding habits of mullet (Mugil Cephalus), milkfish (Chanos Chanos), and the ten-pounder (Elops Machnata). Trans. Am. Fish. Soc. 250–261. doi:10.1577/1548-8659(1944)74[250:fatfci]2.0.co;2

Hillebrand, H., E. Acevedo-Trejos, S. D. Moorthi, A. Ryabov, M. Striebel, P. K. Thomas, and M. Schneider. 2022. Cell size as driver and sentinel of phytoplankton community structure and functioning. Funct. Ecol. 36: 276–293. doi:10.1111/1365-2435.13986

Hoover, D. J., and F. T. Mackenzie. 2009. Fluvial fluxes of water, suspended particulate matter, and nutrients and potential impacts on tropical coastal water biogeochemistry: Oahu, Hawai‘i. Aquat. Geochem. 15: 547–570. doi:10.1007/s10498-009-9067-2

Hothorn, T., F. Bretz, and P. Westfall. 2008. Simultaneous inference in general parametric models. Biometrical Journal 50: 346–363. doi:10.1002/bimj.200810425

Hunter-Cevera, K. R., M. G. Neubert, R. J. Olson, A. R. Solow, A. Shalapyonok, and H. M. Sosik. 2016. Physiological and ecological drivers of early spring blooms of a coastal phytoplankter. Science 354: 326–329. doi:10.1126/science.aaf8536

Hyatt, D., G.-L. Chen, P. F. Locascio, M. L. Land, F. W. Larimer, and L. J. Hauser. 2010. Prodigal: prokaryotic gene recognition and translation initiation site identification. BMC Bioinformatics 11: 119. doi:10.1186/1471-2105-11-119

Inomura, K., J. J. P. Karlusich, S. Dutkiewicz, C. Deutsch, P. J. Harrison, and C. Bowler. 2023. High growth rate of diatoms explained by reduced carbon requirement and low energy cost of silica deposition. Microbiol. Spectr. 11: e03311–22. doi:10.1128/spectrum.03311-22

Irwin, A. J., and M. J. Oliver. 2009. Are ocean deserts getting larger? Geophys. Res. Lett. 36. doi:10.1029/2009gl039883

James, A. K. and others. 2020. An island mass effect resolved near Moʻorea, French Polynesia. Front. Mar. Sci. 7: 368. doi:10.3389/fmars.2020.00016

James, C. C. and others. 2022. Influence of nutrient supply on plankton microbiome biodiversity and distribution in a coastal upwelling region. Nat Commun 13: 2448. doi:10.1038/s41467-022-30139-4

Jokiel. 1991. Jokiel’s illustrated scientific guide to Kāneʻohe Bay. 1–66. doi:10.13140/2.1.3051.9360

Jovel, J. and others. 2016. Characterization of the gut microbiome using 16S or shotgun metagenomics. Front. Microbiol. 7: 459. doi:10.3389/fmicb.2016.00459

Karl, D. M., and M. J. Church. 2014. Microbial oceanography and the Hawaii Ocean Time-series programme. Nature Reviews Microbiology 12: 699–713. doi:10.1038/nrmicro3333

Karl, D. M., and R. Lukas. 1996. The Hawaii Ocean Time-series (HOT) program: Background, rationale and field implementation. Deep Sea Res Part Ii Top Stud Oceanogr 43: 129–156. doi:10.1016/0967-0645(96)00005-7

Kelly, M. 1973. Some legendary and historical aspects of Heeia Fishpond, Koolau, Oahu, Bishop Museum.

Kolde, R. 2019. pheatmap: Pretty Heatmaps. R package version 1.0.12.

Koumandou, V. L., R. E. R. Nisbet, A. C. Barbrook, and C. J. Howe. 2004. Dinoflagellate chloroplasts – where have all the genes gone? Trends Genet. 20: 261–267. doi:10.1016/j.tig.2004.03.008

Kwiatkowski, L., O. Aumont, and L. Bopp. 2018. Consistent trophic amplification of marine biomass declines under climate change. Glob Change Biol 25: 218–229. doi:10.1111/gcb.14468

Kwon, E. Y., M. G. Sreeush, A. Timmermann, D. M. Karl, M. J. Church, S.-S. Lee, and R. Yamaguchi. 2022. Nutrient uptake plasticity in phytoplankton sustains future ocean net primary production. Sci. Adv. 8: eadd2475. doi:10.1126/sciadv.add2475

Lahti, and S. Shetty. 2017. Tools for microbiome analysis in R.

Langmead, B., and S. L. Salzberg. 2012. Fast gapped-read alignment with Bowtie 2. Nat Meth **9**: 357–359. doi:10.1038/nmeth.1923

Laws, E. A. 1975. The importance of respiration losses in controlling the size distribution of marine phytoplankton. Ecology 56: 419–426. doi:10.2307/1934972

Laws, E. A., and C. B. Allen. 1996. Water quality in a subtropical embayment more than a decade after diversion of sewage discharges. Pacific Science 50: 194–210.

Laws, E. A., and D. G. Redalje. 1982. Sewage diversion effects on the water column of a subtropical estuary. Mar. Environ. Res. 6: 265–279. doi:10.1016/0141-1136(82)90041-1

Laws, E., and S. Taguchi. 2018. The 1987–1989 phytoplankton bloom in Kaneohe Bay. Water 10: 747–9. doi:10.3390/w10060747

Lee, M. D. 2019. GToTree: a user-friendly workflow for phylogenomics. Bioinformatics 35: btz188. doi:10.1093/bioinformatics/btz188

Lee, M. D., N. A. Ahlgren, J. D. Kling, N. G. Walworth, G. Rocap, M. A. Saito, D. A. Hutchins, and E. A. Webb. 2019. Marine Synechococcus isolates representing globally abundant genomic lineages demonstrate a unique evolutionary path of genome reduction without a decrease in GC content. Environmental Microbiology. doi:10.1111/1462-2920.14552

Leng, Q., X. Guo, J. Zhu, and A. Morimoto. 2023. Contribution of the open ocean to the nutrient and phytoplankton inventory in a semi-enclosed coastal sea. Biogeosciences 20: 4323–4338. doi:10.5194/bg-20-4323-2023

Li, B., D. M. Karl, R. M. Letelier, R. R. Bidigare, and M. J. Church. 2013. Variability of chromophytic phytoplankton in the North Pacific Subtropical Gyre. Deep Sea Res. Part II: Top. Stud. Oceanogr. 93: 84–95. doi:10.1016/j.dsr2.2013.03.007

Li, F., J. Beardall, K. Gao, and H. editor: S. Sathyendranath. 2018. Diatom performance in a future ocean: interactions between nitrogen limitation, temperature, and CO2-induced seawater acidification. ICES J. Mar. Sci. 75: 1451–1464. doi:10.1093/icesjms/fsx239

Li, G., L. Cheng, J. Zhu, K. E. Trenberth, M. E. Mann, and J. P. Abraham. 2020. Increasing ocean stratification over the past half-century. Nat Clim Change 10: 1116–1123. doi:10.1038/s41558-020-00918-2

Lin, H., and S. D. Peddada. 2024. Multigroup analysis of compositions of microbiomes with covariate adjustments and repeated measures. Nat. Methods 21: 83–91. doi:10.1038/s41592-023-02092-7

Lowe, R. J., J. L. Falter, S. G. Monismith, and M. J. Atkinson. 2009. A numerical study of circulation in a coastal reef-lagoon system. Journal of Geophysical Research 114: 997. doi:10.1029/2008jc005081

Mantoura, R. F. C., and C. A. Llewellyn. 1983. The rapid determination of algal chlorophyll and carotenoid pigments and their breakdown products in natural waters by reverse-phase high-performance liquid chromatography. Anal Chim Acta 151: 297–314. doi:10.1016/s0003-2670(00)80092-6

Martiny, A. C. and others. 2022. Marine phytoplankton resilience may moderate oligotrophic ecosystem responses and biogeochemical feedbacks to climate change. Limnol. Oceanogr. 67: S378–S389. doi:10.1002/lno.12029

McKenzie, T., H. Dulai, and J. Chang. 2019. Parallels between stream and coastal water quality associated with groundwater discharge. PLoS ONE 14: e0224513. doi:10.1371/journal.pone.0224513

McKenzie, T., S. Habel, and H. Dulai. 2021. Sea-level rise drives wastewater leakage to coastal waters and storm drains. Limnol. Oceanogr. Lett. 6: 154–163. doi:10.1002/lol2.10186

McMurdie, P. J., and S. Holmes. 2013. phyloseq: an R package for reproducible interactive analysis and graphics of microbiome census data. M. Watson [ed.]. PLoS ONE 8: e61217. doi:10.1371/journal.pone.0061217

Mende, D. R., J. A. Bryant, F. O. Aylward, J. M. Eppley, T. Nielsen, D. M. Karl, and E. F. DeLong. 2017. Environmental drivers of a microbial genomic transition zone in the ocean’s interior. Nat Microbiol 2: 1367–1373. doi:10.1038/s41564-017-0008-3

Messié, M., A. Petrenko, A. M. Doglioli, C. Aldebert, E. Martinez, G. Koenig, S. Bonnet, and T. Moutin. 2020. The delayed island mass effect: how islands can remotely trigger blooms in the oligotrophic ocean. Geophys Res Lett 47. doi:10.1029/2019gl085282

Messié, M., A. Petrenko, A. M. Doglioli, E. Martinez, and S. Alvain. 2022. Basin-scale biogeochemical and ecological impacts of islands in the tropical Pacific Ocean. Nat Geosci 1–6. doi:10.1038/s41561-022-00957-8

Minoche, A. E., J. C. Dohm, and H. Himmelbauer. 2011. Evaluation of genomic high-throughput sequencing data generated on Illumina HiSeq and Genome Analyzer systems. Genome Biol 12: R112. doi:10.1186/gb-2011-12-11-r112

Möhlenkamp, P., C. Beebe, M. McManus, A. Kawelo, K. Kotubetey, M. Lopez-Guzman, C. Nelson, and R. Alegado. 2019. Kū Hou Kuapā: Cultural restoration improves water budget and water quality dynamics in Heʻeia Fishpond. Sustainability 11: 161–25. doi:10.3390/su11010161

Monger, B. C., and M. R. Landry. 1993. Flow cytometric analysis of marine bacteria with Hoechst 33342. Applied and Environmental Microbiology 59: 905–911. doi:10.1128/aem.59.3.905-911.1993

Moore, J. K. and others. 2018. Sustained climate warming drives declining marine biological productivity. Science 359: 1139–1143. doi:10.1126/science.aao6379

Nuʻuhiwa, K. 2019. Papakū makawalu: A Methodology and Pedagogy of Understanding the Hawaiian Universe, p. 39–49. *In* N. Wilson-Hokowhitu [ed.], The Past Before Us : Mookauhau As Methodology. University of Hawaii Press. 10.1515/9780824878177-007

Parada, A. E., D. M. Needham, and J. A. Fuhrman. 2016. Every base matters: assessing small subunit rRNA primers for marine microbiomes with mock communities, time series and global field samples. Environmental Microbiology 18: 1403–1414. doi:10.1111/1462-2920.13023

Pasulka, A. L., M. R. Landry, D. A. A. Taniguchi, A. G. Taylor, and M. J. Church. 2013. Temporal dynamics of phytoplankton and heterotrophic protists at station ALOHA. Deep Sea Res Part Ii Top Stud Oceanogr 93: 44–57. doi:10.1016/j.dsr2.2013.01.007

Pritchard, L., R. H. Glover, S. Humphris, J. G. Elphinstone, and I. K. Toth. 2015. Genomics and taxonomy in diagnostics for food security: soft-rotting enterobacterial plant pathogens. Anal. Methods 8: 12–24. doi:10.1039/c5ay02550h

Quast, C., E. Pruesse, P. Yilmaz, J. Gerken, T. Schweer, P. Yarza, J. Peplies, and F. O. Glöckner. 2012. The SILVA ribosomal RNA gene database project: improved data processing and web-based tools. Nucleic Acids Res 41: D590–D596. doi:10.1093/nar/gks1219

R Core Team. 2021. R: A language and environment for statistical computing. R Foundation for Statistical Computing, Vienna, Austria. https://www.R-project.org/.

Redfield, A. C. 1960. The biological control of chemical factors in the environment. Sci Prog 11: 150–70. doi:10.1086/646891

Ribalet, F. and others. 2010. Unveiling a phytoplankton hotspot at a narrow boundary between coastal and offshore waters. Proc National Acad Sci 107: 16571–16576. doi:10.1073/pnas.1005638107

Rii, Y. M., S. Duhamel, R. R. Bidigare, D. M. Karl, D. J. Repeta, and M. J. Church. 2016. Diversity and productivity of photosynthetic picoeukaryotes in biogeochemically distinct regions of the South East Pacific Ocean. Limnol. Oceanogr. 61: 806–824. doi:10.1002/lno.10255

Ringuet, S., and F. T. Mackenzie. 2005. Controls on nutrient and phytoplankton dynamics during normal flow and storm runoff conditions, southern Kaneohe Bay, Hawaii. Estuaries 28: 327–337. doi:10.1007/bf02693916

Rocap, G., D. L. Distel, J. B. Waterbury, and S. W. Chisholm. 2002. Resolution of Prochlorococcus and Synechococcus ecotypes by using 16S-23S ribosomal DNA internal transcribed spacer sequences. Appl. Environ. Microbiol. 68: 1180–1191. doi:10.1128/aem.68.3.1180-1191.2002

Ruf, T. 1999. The Lomb-Scargle Periodogram in biological rhythm research: analysis of incomplete and unequally spaced time-series. Biol Rhythm Res 30: 178–201. doi:10.1076/brhm.30.2.178.1422

Schaum, C.-E. and others. 2017. Adaptation of phytoplankton to a decade of experimental warming linked to increased photosynthesis. Nat. Ecol. Evol. 1: 0094. doi:10.1038/s41559-017-0094

Selph, K., E. Goetze, M. Jungbluth, P. Lenz, and G. Kolker. 2018. Microbial food web connections and rates in a subtropical embayment. Mar Ecol Prog Ser 590: 19–34. doi:10.3354/meps12432

Smith, S. V. 1981. Responses of Kaneohe Bay, Hawaii, to relaxation of sewage stress, p. 391–410. *In* Neilson, B.J., Cronin, L.E. (eds) Estuaries and Nutrients. Contemporary Issues in Science and Society. Humana Press. 10.1007/978-1-4612-5826-1_18

Stock, C. A. and others. 2017. Reconciling fisheries catch and ocean productivity. Proc National Acad Sci 114: E1441–E1449. doi:10.1073/pnas.1610238114

Stoecker, D. K., P. J. Hansen, D. A. Caron, and A. Mitra. 2016. Mixotrophy in the marine plankton. Annu Rev Mar Sci 9: 1–25. doi:10.1146/annurev-marine-010816-060617

Strimmer, K. 2008. fdrtool: a versatile R package for estimating local and tail area-based false discovery rates. Bioinformatics 24: 1461–1462. doi:10.1093/bioinformatics/btn209

Stukel, M. R., J. P. Irving, T. B. Kelly, M. D. Ohman, C. K. Fender, and N. Yingling. 2023. Carbon sequestration by multiple biological pump pathways in a coastal upwelling biome. Nat. Commun. 14: 2024. doi:10.1038/s41467-023-37771-8

Suzuki, M. T., O. Béjà, L. T. Taylor, and E. F. DeLong. 2001. Phylogenetic analysis of ribosomal RNA operons from uncultivated coastal marine bacterioplankton. Environmental Microbiology 3: 323–331. doi:10.1046/j.1462-2920.2001.00198.x

Tucker, S. J., K. C. Freel, E. A. Monaghan, C. E. S. Sullivan, O. Ramfelt, Y. M. Rii, and M. S. Rappé. 2021. Spatial and temporal dynamics of SAR11 marine bacteria across a nearshore to offshore transect in the tropical Pacific Ocean. Peerj 9: e12274. doi:10.7717/peerj.12274

Ustick, L. J., A. A. Larkin, and A. C. Martiny. 2023. Global scale phylogeography of functional traits and microdiversity in Prochlorococcus. ISME J. 17: 1671–1679. doi:10.1038/s41396-023-01469-y

Utter, D. R., G. G. Borisy, A. M. Eren, C. M. Cavanaugh, and J. L. M. Welch. 2020. Metapangenomics of the oral microbiome provides insights into habitat adaptation and cultivar diversity. 1: 808410–40. doi:10.1101/2020.05.01.072496

Vollbrecht, C., P. Moehlenkamp, J. M. Gove, A. B. Neuheimer, and M. A. McManus. 2021. Long-term presence of the island mass effect at Rangiroa Atoll, French Polynesia. Frontiers Mar Sci 7: 595294. doi:10.3389/fmars.2020.595294

Vrana, I. and others. 2023. Successful acclimation of marine diatoms Chaetoceros curvisetus/pseudocurvisetus to climate change. Limnol. Oceanogr. 68: S158–S173. doi:10.1002/lno.12293

Ward, B. A., and M. J. Follows. 2016. Marine mixotrophy increases trophic transfer efficiency, mean organism size, and vertical carbon flux. Proc. Natl. Acad. Sci. 113: 2958–2963. doi:10.1073/pnas.1517118113

Wei, T., and V. Simko. 2021. corrplot: A visualization of a correlation matrix.

Welschmeyer, N. A. 1994. Fluorometric analysis of chlorophyll a in the presence of chlorophyll b and pheopigments. Limnol. Oceanogr. 39: 1985–1992. doi:10.4319/lo.1994.39.8.1985

Wickham, H. 2016. ggplot2: elegant graphics for data analysis. Springer-Verlag New York.

Williams, I. D., J. K. Baum, A. Heenan, K. M. Hanson, M. O. Nadon, and R. E. Brainard. 2015. Human, oceanographic and habitat drivers of central and western Pacific coral reef fish assemblages. Plos One 10: e0120516. doi:10.1371/journal.pone.0120516

Willis, A. D., and B. D. Martin. 2020. Estimating diversity in networked ecological communities. Biostatistics 23: kxaa015. doi:10.1093/biostatistics/kxaa015

Winter, K. and others. 2020a. Ecomimicry in Indigenous resource management: optimizing ecosystem services to achieve resource abundance, with examples from Hawaiʻi. Ecol. Soc. 25. doi:10.5751/es-11539-250226

Winter, K. and others. 2020b. Collaborative research to inform adaptive comanagement: a framework for the Heʻeia National Estuarine Research Reserve. Ecol Soc 25. doi:10.5751/es-11895-250415

Wright, E. 2016. Using DECIPHER v2.0 to analyze big biological sequence data in R. The R Journal.

Yamaguchi, R., and T. Suga. 2019. Trend and variability in global upper-ocean stratification since the 1960s. J. Geophys. Res.: Oceans 124: 8933–8948. doi:10.1029/2019jc015439

Yan, W. and others. 2022. Diverse subclade differentiation attributed to the ubiquity of Prochlorococcus High-Light-Adapted Clade II. Mbio 13: e03027–21. doi:10.1128/mbio.03027-21

Yeh, Y., J. McNichol, D. M. Needham, E. B. Fichot, L. Berdjeb, and J. A. Fuhrman. 2021. Comprehensive single-PCR 16S and 18S rRNA community analysis validated with mock communities, and estimation of sequencing bias against 18S. Environ Microbiol 23: 3240– 3250. doi:10.1111/1462-2920.15553

Yeo, S. K., M. J. Huggett, A. Eiler, and M. S. Rappé. 2013. Coastal bacterioplankton community dynamics in response to a natural disturbance D.L. Kirchman [ed.]. PLoS ONE 8: e56207–14. doi:10.1371/journal.pone.0056207

Zhang, C., Y. Wang, K. Hamilton, and A. Lauer. 2016. Dynamical downscaling of the climate for the Hawaiian Islands. Part II: Projection for the Late Twenty-First Century. J. Clim. 29: 8333–8354. doi:10.1175/jcli-d-16-0038.1

