## Supplemental Information for "Seasonal and spatial transitions in phytoplankton assemblages spanning estuarine to open ocean waters of the tropical Pacific"

Keliʻiahonui Kotubetey[4]

A. Hiʻilei Kawelo[4]

Kawika B. Winter[1,3] https://orcid.org/0000-0003-3762-7125

Michael S. Rappé[1] https://orcid.org/0000-0002-9829-251X

[1]Hawaiʻi Institute of Marine Biology, University of Hawaiʻi at Mānoa, Kāneʻohe, Hawaiʻi, USA, 96744

[2]Marine Biology Graduate Program, University of Hawaiʻi at Mānoa, Honolulu, Hawaiʻi, USA, 96822

[3]Heʻeia National Estuarine Research Reserve, Kāneʻohe, Hawaiʻi, USA, 96744

[4]Paepae o Heʻeia, Kāneʻohe Hawaiʻi, USA, 96744

[#]Josephine Bay Paul Center for Comparative Molecular Biology and Evolution, Marine Biological Laboratory, Woods Hole, MA 02543, USA

**Running Title:** Spatiotemporal shifts in phytoplankton

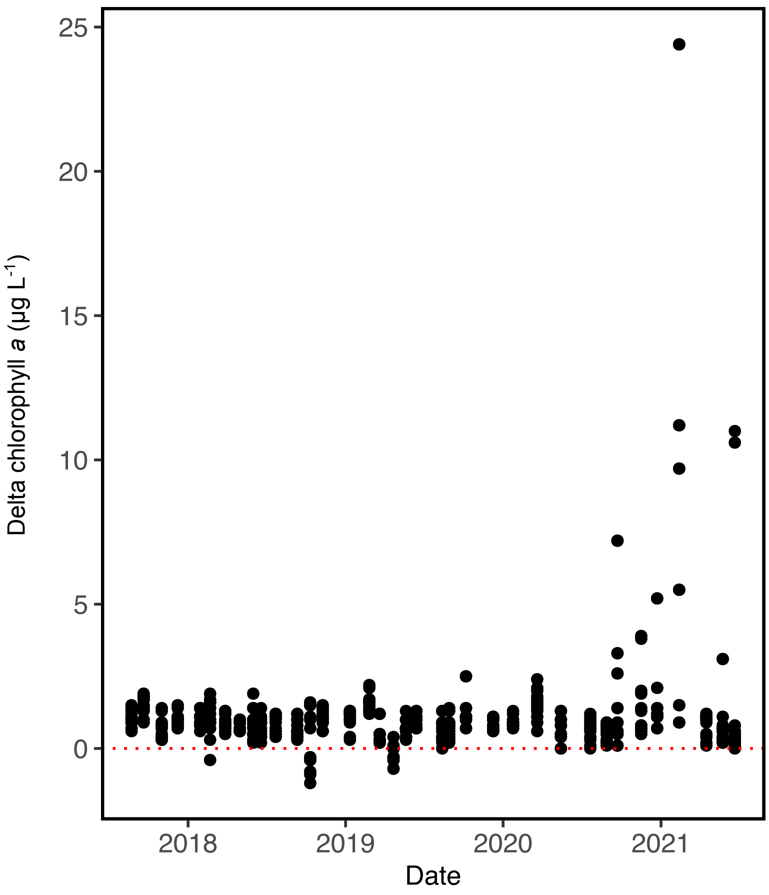


**Fig. S1.** Persistence of chlorophyll *a* enhancement. Delta chlorophyll *a* calculated as the chlorophyll *a* concentrations measured at each station in the estuarine He‘eia Fishpond and coastal Kāne‘ohe Bay (Wai2, Kaho‘okele, HP1, AR, SB, SR8, NB,CB) minus the chlorophyll *a* concentrations measured at stations positioned furthest offshore (STO1, NTO1) during the same sampling event.

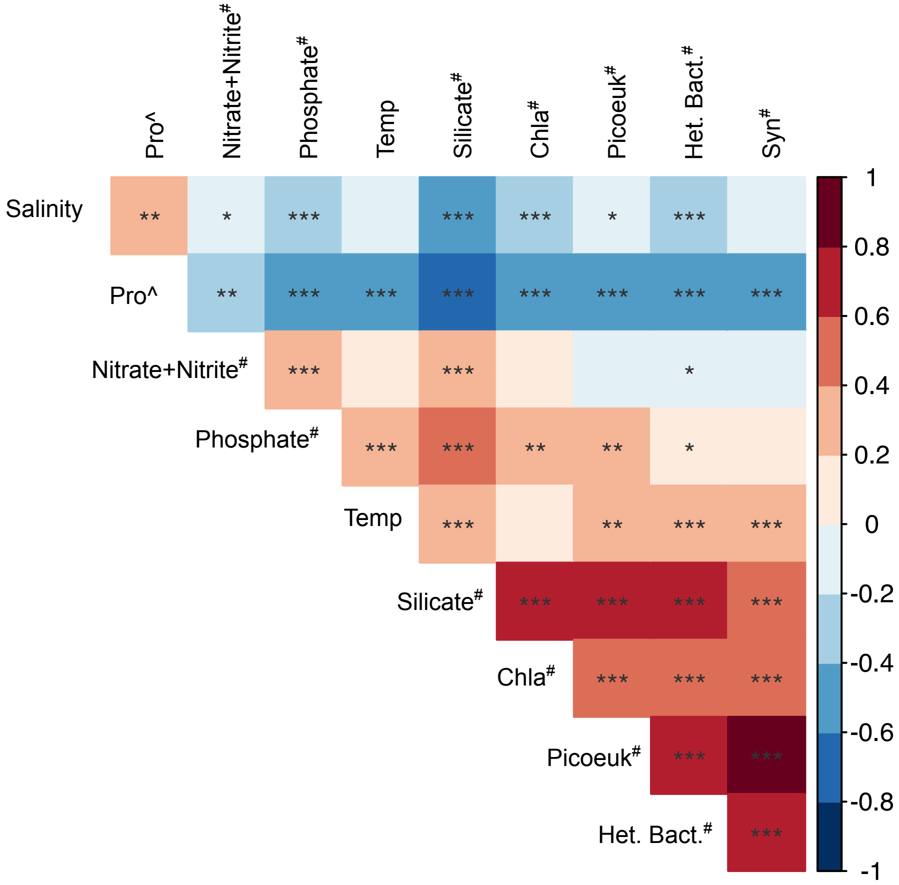


**Fig. S2.** Pearson's correlation of biogeochemical parameters across all KByT stations.
A pound symbol (#) denotes variables with log transformations, while a carrot (^) denotes variables with log+1 transformations. Asterisks represent significant correlations ( *, p<0.05; **, p<0.01; ***, p<0.001).
Abbreviations and Units: Het.Bac: heterotrophic bacteria (cells $mL^{-1}$); Syn: *Synechococcus* (cells $mL^{-1}$);
Pro: *Prochlorococcus* (cells $mL^{-1}$); Picoeuk: Photosynthetic picoeukaryotes (cells $mL^{-1}$);
Temp: Seawater temperature (°C); Chla: Chlorophyll a (μg $L^{-1}$); Salinity (ppt); Phosphate, Nitrate+Nitrite, Silicate (μM).

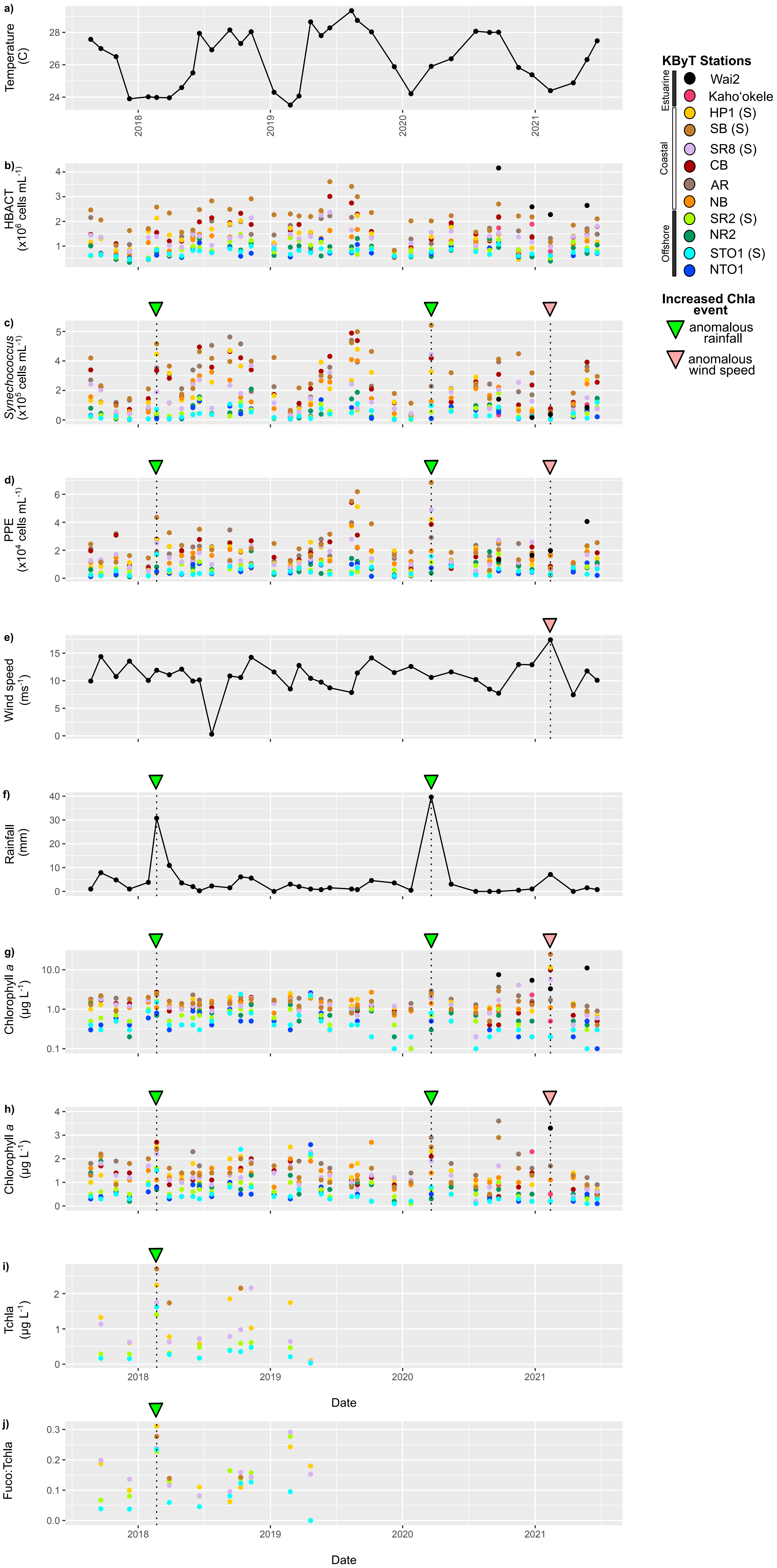


**Fig. S3**. Seasonal changes in sea surface temperature (°C; measured at station HP1) aligns with changes in in the cellular abundances of heterotrophic bacteria (HBACT), *Synechococcus*, and photosynthetic picoeukaryotes (PPE) (a,b,c,d). Three sampling events (February 21, 2018, March 20, 2020, and February 12, 2021) show high rainfall or winds speeds and coincide with elevated chlorophyll *a* concentrations (e,f,g,h,i). Changes in chlorophyll *a* concentrations (Chla) over time for the entire range of chlorophyll *a* concentrations (g), a zoom in range displaying just chlorophyll *a* concentrations between 0-4 µg $L^{-1}$ (h), and total chlorophyll *a* (Tchla) measured via high-performance liquid chromatography (HPLC). Shifts in the cellular abundances of *Synechococcus* and photosynthetic picoeukaryotes are observed during the three episodic storm events. Increases in the ratio of fucoxanthin:total chlorophyll *a* (Fuco:Tchla), a pigment indicative of diatoms, is elevated during the February 21, 2018 storm event for stations in southern Kāne'ohe Bay and adjacent offshore, suggesting some storm events cause phytoplankton responses at relatively large spatial scales. Stations in southern Kāne'ohe Bay and the adjacent offshore with HPLC data are denoted with an (S) in the key.

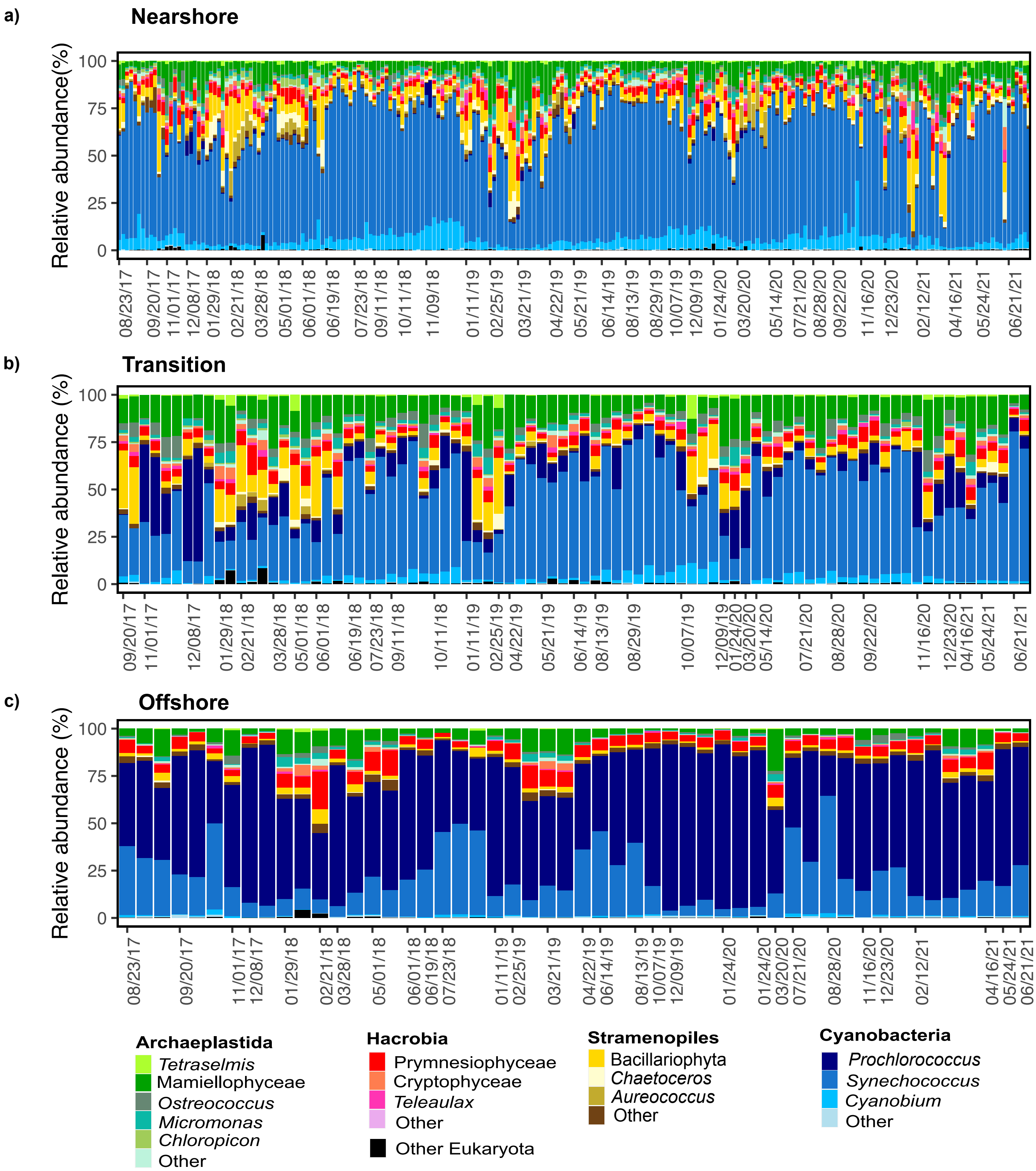


**Fig. S4.** The relative abundance of major phytoplankton groups differs across the three community types: **a)** nearshore, **b)** transition, and **c)** offshore. Phytoplankton classes and genera within Archaeplastida, Hacrobia, Stramenopiles, and Cyanobacteria with >1% average relative abundance in at least one community type are specified. Phytoplankon from Archaeplastida, Hacrobia, Stramenopiles, and Cyanobacteria that are below this threshold are assigned as Other within their respective supergroup. Rhizaria, Excavata, and eukaryotes unclassified below the domain level were assinged to Other Eukaryota. Within each community type samples are grouped first by sampling event (36 total between 2017-2021), then by the station.

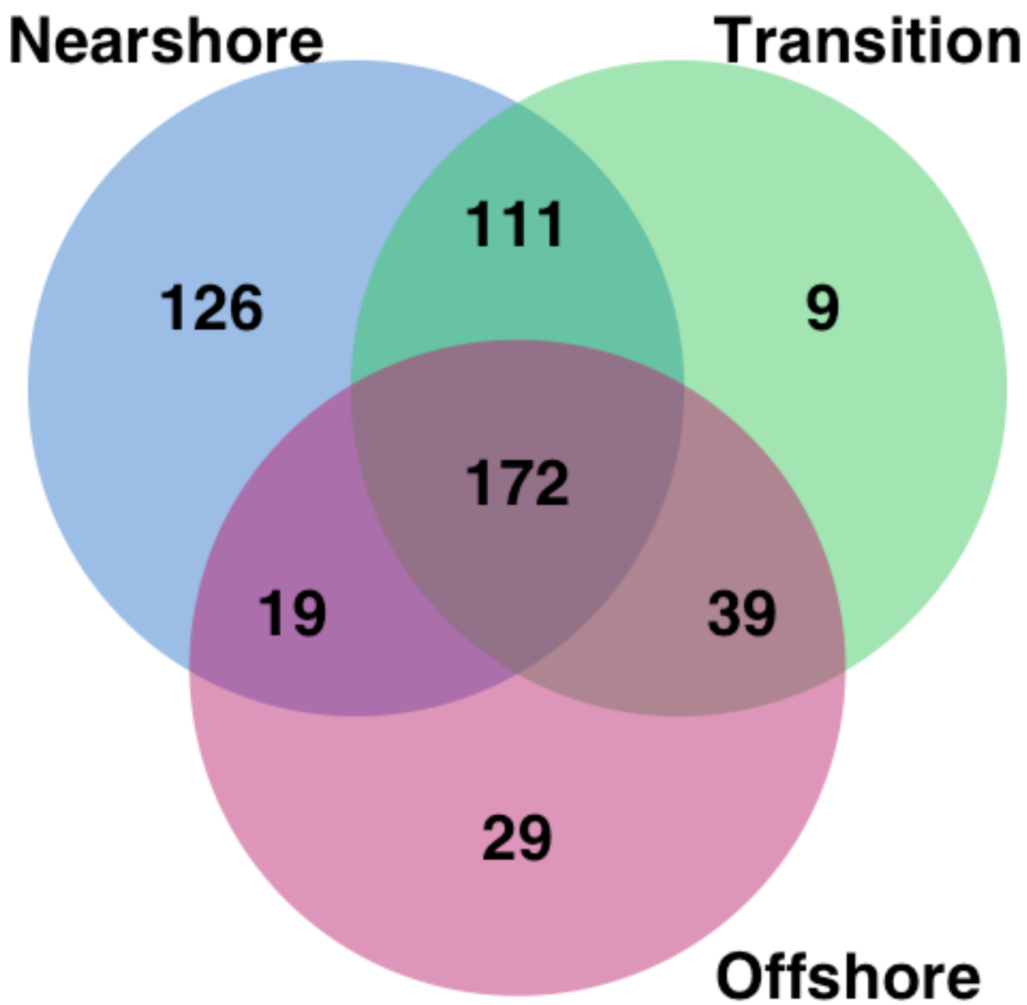


**Fig. S5**. Count of ASVs shared and unique to the three community types.

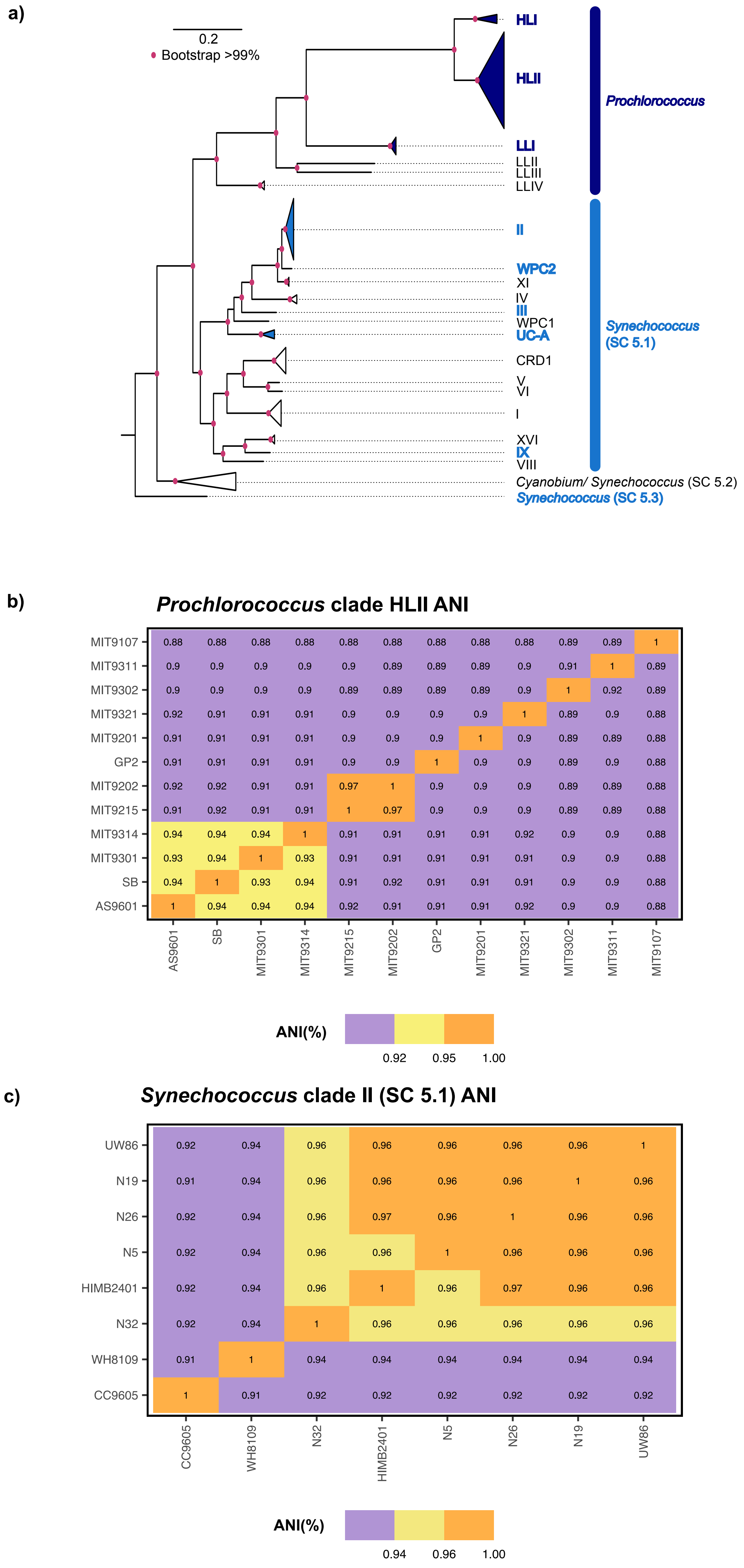


**Fig. S6**. **a**) Phylogenomic tree based on cyanobacterial marker genes found in 56 Prochlorococcus, Synechococcus, and Cyanobium isolate genomes and outgroup (Gloeobacter violaceus- not shown). Clades detected in surface ocean metagenomic samples from the Kāneʻohe Bay Time-series (KByT) and Station ALOHA are colored and bolded.Average nucleotide identity (ANI) between closely related Prochlorococcus Clade HLII isolate genomes (**b**),as well as those for Synechococcus Clade II isolate genomes (**c**).

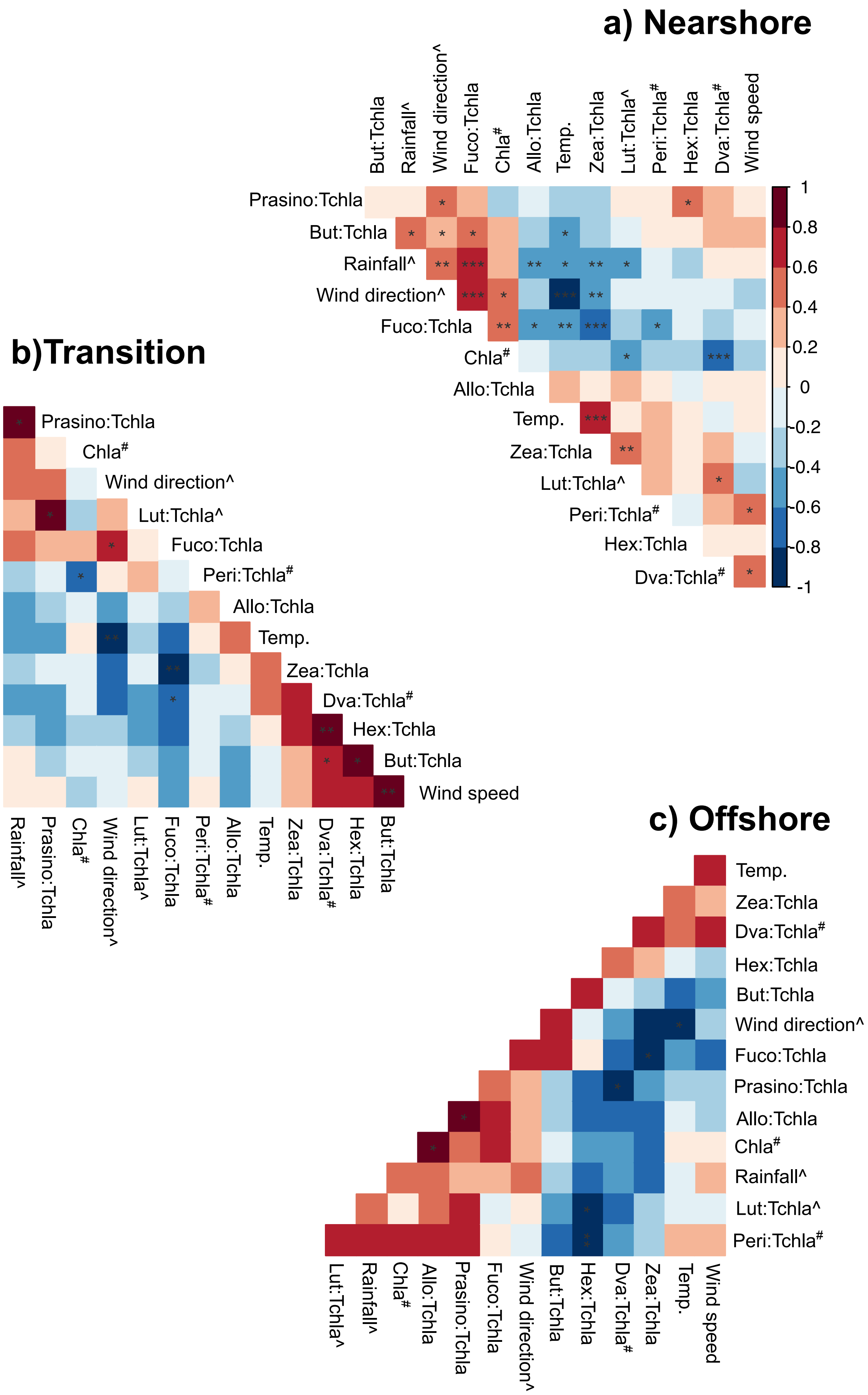


**Fig. S7**. Pearson's correlation between phytoplankton pigment: total chlorophyll *a* (Tchla) ratios and seawater temperature and weather parameters in the **a**) nearshore, **b**) transition, and **c**) offshore community types. A pound symbol (#) denotes variables with log transformations, while a carrot (^) denotes variables with log+1 transformations. Asterisks represent significant correlations ( *, p<0.05; **, p<0.01; ***, p<0.001). Abbreviations: Fuco: Fucoxanthin, Peri: Peridinin; Allo: Alloxanthin; Prasino: Prasinoxanthin; Hex: 19′-hexanoyloxyfucoxanthin; But: 19′-butanoyloxyfucoxanthin; Zea: zeaxanthin; DVchla: divinyl chlorophyll *a*; Chla: fluorometric chlorophyll *a* (µg $L^{-1}$), Temp: Seawater temperature (°C); Wind direction (degrees); Wind Speed ($ms^{-1}$); Rainfall (mm).

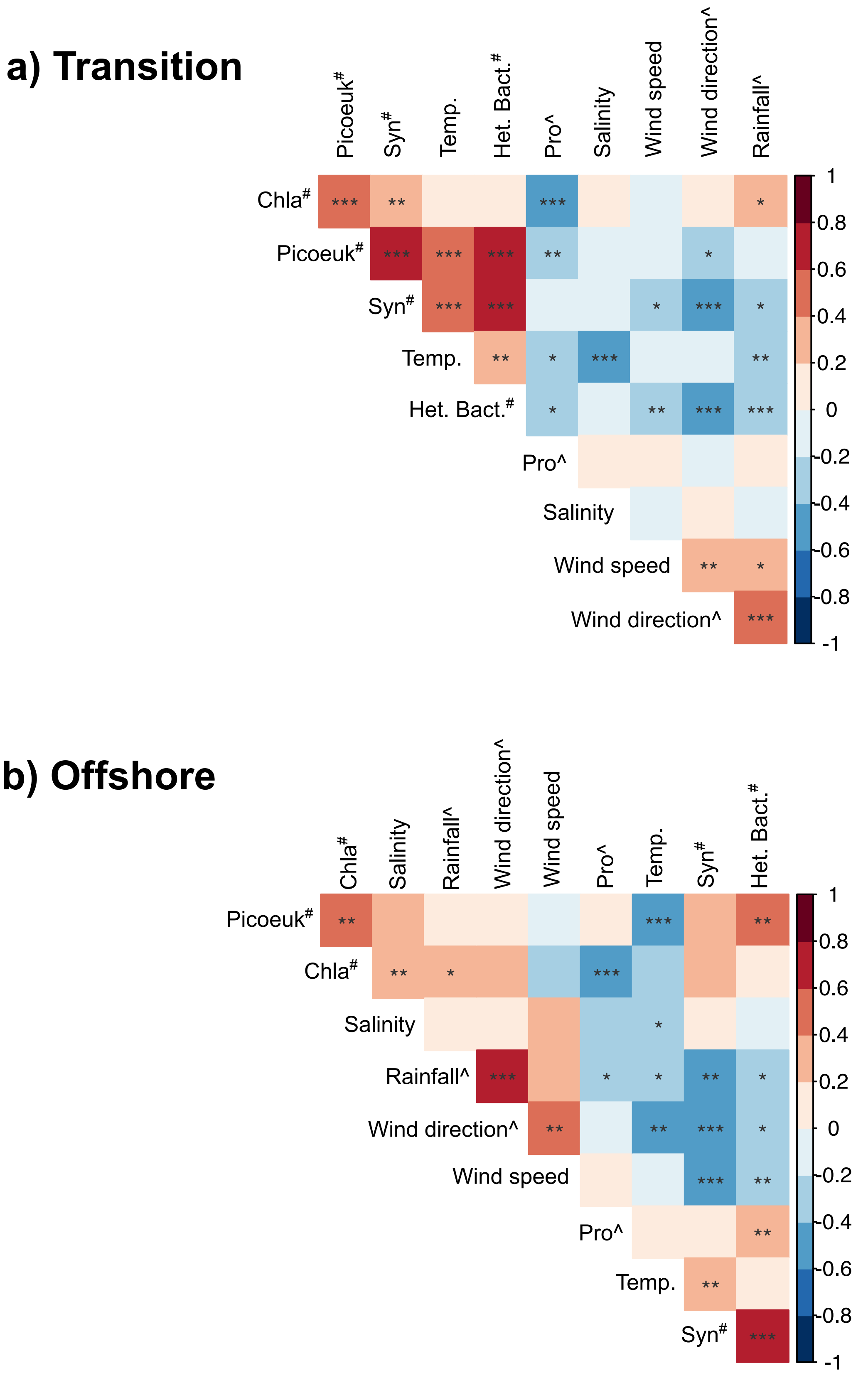


**Fig. S8.** Pearson's correlation across biogeochemical parameters in the **a)** transition and **b)** offshore community type. A pound symbols (#) denotes variables with log transformations, while a carrot (^) denotes variables with log+1 transformations. Asterisks represent significant correlations ( *, p<0.05; **, p<0.01; ***, p<0.001). Abbreviations and Units: Het.Bac: heterotrophic bacteria (cells $mL^{-1}$); Syn: *Synechococcus* (cells $mL^{-1}$); Pro: *Prochlorococcus* (cells $mL^{-1}$); Picoeuk: Photosynthetic picoeukaryotes (cells $mL^{-1}$); Temp: Seawater temperature (°C); Chla: Chlorophyll a (μg $L^{-1}$); Salinity (ppt); Phosphate, Nitrate+Nitrite, Silicate (μM); Wind direction (degrees); Wind Speed ($ms^{-1}$); Rainfall (mm).

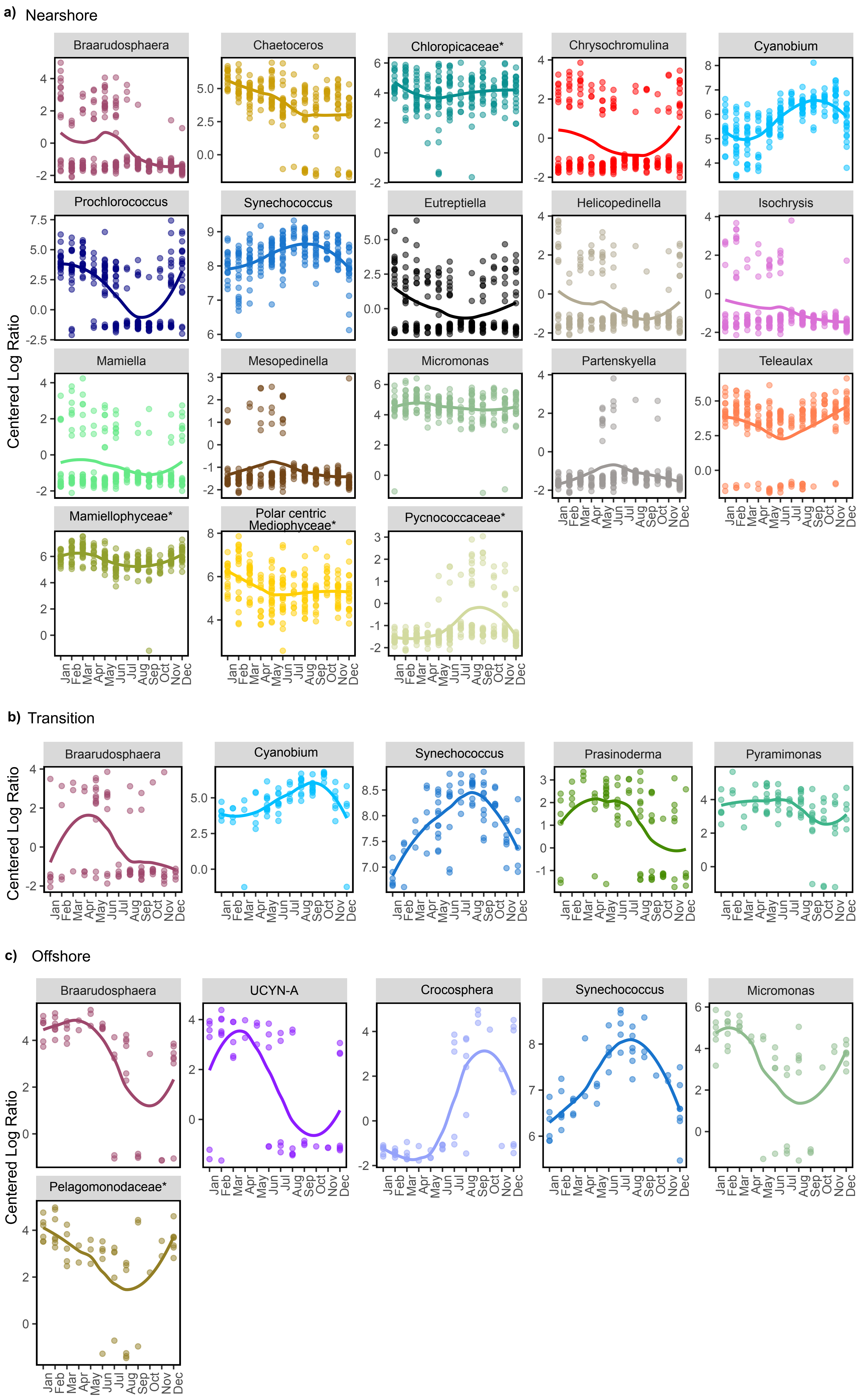


**Fig. S9.** Phytoplankton genera with signficant seasonality in each community type as defined by Lomb-Scargle Periodicity test. Phytoplankton groups that were classified at the family-level but unidentified at the genus-level are denoted with an asterisk. A local polynomial regression fit line is shown for each genus.

**Table S1.** Biogeochemical parameters across stations. Abbreviations: DIN:P: Dissolved inorganic nitrogen to phosphate.

| | | WAI2 | KAHO | HP1 | SB | SR8 | AR | CB | NB | SR2 | NR2 | STO1 | NTO1 | ALOHA |
|---|---|---|---|---|---|---|---|---|---|---|---|---|---|---|
| **Latitude** | | 21.43833 N | 21.43582 N | 21.43688 N | 21.43635 N | 21.44983333 N | 21.46765 N | 21.45705 N | 21.49048333 N | 21.4691667 N | 21.5147 N | 21.4829 N | 21.526 N | 22.45 N |
| **Longitude** | | 157.81044 W | 157.80546 W | 157.80331 W | 157.7773667 W | 157.7932167 W | 157.83585 W | 157.81135 W | 157.8332833 W | 157.77752 W | 157.8117167 W | 157.7663 W | 157.7979 W | 158.00 W |
| **Seawater temp. (ºC)** | n | 4 | 4 | 36 | 36 | 36 | 36 | 36 | 35 | 35 | 36 | 36 | 36 | 95 |
| | mean±sd | 28.3±1.7 | 27±1.1 | 26.4±1.7 | 26±1.8 | 26.1±1.5 | 26.4±2.0 | 26.2±1.8 | 26.2±1.7 | 25.6±1.3 | 25.9±1.5 | 25.6±1.3 | 25.7±1.3 | 26.0±1.4 |
| **pH** | n | 4 | 4 | 36 | 36 | 36 | 36 | 36 | 35 | 35 | 36 | 36 | 36 | 28 |
| | mean±sd | 7.8±0.3 | 7.9±0.5 | 7.7±0.3 | 7.9±0.2 | 7.9±0.1 | 7.9±0.1 | 7.9±0.2 | 7.9±0.1 | 7.8±0.2 | 8.0±0.2 | 8.0±0.2 | 8.0±0.1 | 8.1±0 |
| **Salinity** | n | 4 | 4 | 36 | 36 | 36 | 36 | 36 | 35 | 35 | 36 | 36 | 36 | 95 |
| | mean±sd | 30±0.7 | 34.1±0.6 | 34.0±1.1 | 34.3±1.1 | 34.3±1.1 | 33.9±1.2 | 34.3±1.4 | 34.3±1 | 34.6±1.1 | 34.7±1 | 34.7±1.1 | 34.7±1 | 34.8±0.2 |
| **Chlorophyll a (µg L$^{-1}$)** | n | 4 | 4 | 36 | 36 | 36 | 36 | 36 | 35 | 35 | 36 | 36 | 36 | 54 |
| | mean±sd | 6.8±3.3 | 1.0±0.9 | 1.7±1.7 | 2.1±3.9 | 1.4±1 | 1.8±0.5 | 1.5±1.5 | 1.3±0.4 | 0.7±0.4 | 0.7±0.4 | 0.5±0.5 | 0.5±0.4 | 0.09±0.03 |
| **Photosynthetic picoeukaryotes (cells mL$^{-1}$)*10$^{4}$** | n | 4 | 4 | 36 | 35 | 36 | 36 | 36 | 35 | 35 | 36 | 36 | 36 | 22 |
| | mean±sd | 2.24±1.24 | 0.73±0.5 | 1.67±1 | 2.37±1.46 | 1.48±0.86 | 1.88±0.76 | 1.90±1 | 1.43±0.67 | 0.71±0.48 | 0.73±0.45 | 0.54±0.36 | 0.53±0.35 | 0.11±0.04 |
| **Heterotrophic bacteria (cells mL$^{-1}$) *10$^{6}$** | n | 4 | 4 | 36 | 35 | 36 | 36 | 36 | 35 | 35 | 36 | 36 | 36 | 22 |
| | mean±sd | 2.92±0.84 | 1.29±0.6 | 1.38±0.44 | 2.16±0.61 | 1.35±0.34 | 1.59±0.37 | 1.59±0.56 | 1.1±0.3 | 0.9±0.25 | 0.82±0.26 | 0.74±0.14 | 0.75±0.19 | 0.49±0.69 |
| **Synechococcus (cells mL$^{-1}$) *10$^{4}$** | n | 4 | 4 | 36 | 35 | 36 | 36 | 36 | 35 | 35 | 36 | 36 | 36 | 22 |
| | mean±sd | 6.97±5.54 | 4.39±4.17 | 18.51±13.33 | 31.07±15.07 | 15.46±10.43 | 24.50±14.61 | 25.38±14.61 | 15.06±10.88 | 4.92±4.12 | 5.55±5.5 | 4.08±3.43 | 4.09±3.82 | 0.19±0.18 |
| **Prochlorococcus (cells mL $^{-1}$) *10$^{4}$** | n | 4 | 4 | 36 | 35 | 36 | 36 | 36 | 35 | 35 | 36 | 36 | 36 | 21 |
| | mean±sd | 0.02±0.01 | 0.04±0.04 | 0.36±0.55 | 0.59±0.69 | 0.73±0.76 | 0.56±0.89 | 0.61±0.8 | 0.54±0.62 | 4.94±4.3 | 3.83±5.81 | 15.70±8.5 | 13.21±7.47 | 20.16±2.68 |
| **Phosphate (µM)** | n | 4 | 4 | 24 | 24 | 24 | 8 | 8 | 8 | 23 | 8 | 24 | 8 | - |
| | mean±sd | 0.36±0.09 | 0.18±0.03 | 0.11±0.04 | 0.09±0.04 | 0.09±0.03 | 0.13±0.07 | 0.09±0.04 | 0.10±0.05 | 0.09±0.02 | 0.09±0.04 | 0.07±0.02 | 0.09±0.02 | - |
| **Silicate (µM)** | n | 4 | 4 | 24 | 24 | 24 | 8 | 8 | 8 | 23 | 8 | 24 | 8 | 51 |
| | mean±sd | 107.33±45.38 | 18.88±7.93 | 11.14±4.66 | 7.21±2.7 | 5.29±3.06 | 18.17±10.31 | 6.56±2.34 | 9.8±2.69 | 2.56±2.03 | 2.25±0.85 | 1.56±0.81 | 1.55±0.44 | 1.23±0.43 |
| **Nitrate+Nitrite (µM)** | n | 4 | 4 | 24 | 24 | 24 | 8 | 8 | 8 | 23 | 8 | 24 | 8 | - |
| | mean±sd | 0.48±0.22 | 1.05±0.6 | 0.44±0.27 | 0.1±0.1 | 0.22±0.16 | 0.32±0.25 | 0.09±0.07 | 0.46±0.37 | 0.28±0.19 | 0.31±0.16 | 0.13±0.12 | 0.09±0.04 | - |
| **Ammonia (µM)** | n | 4 | 4 | 15 | 3 | 6 | 5 | - | 4 | 7 | 2 | 4 | - | - |
| | mean±sd | 0.54±0.52 | 0.98±0.47 | 0.22±0.16 | 0.04±0.01 | 0.04±0.02 | 0.25±0.34 | - | 0.06±0.03 | 0.14±0.19 | 0.06±0.03 | 0.05±0.02 | - | - |
| **DIN:P** | n | 4 | 4 | 15 | 3 | 4 | 5 | - | 4 | 7 | 2 | 4 | - | - |
| | mean±sd | 2.9±2.15 | 11.78±6.47 | 6.85±3.05 | 1.71±0.39 | 11.78±6.47 | 5.89±3.31 | - | 8.31±7.07 | 5.99±3.08 | 6.44±1.14 | 3.02±2.01 | - | - |
| **Low-Level Nitrogen (µM)** | n | - | - | - | - | - | - | - | - | - | - | - | - | 21 |
| | mean±sd | - | - | - | - | - | - | - | - | - | - | - | - | 0.003±0.001 |
| **Low-Level Phosphate (µM)** | n | - | - | - | - | - | - | - | - | - | - | - | - | 29 |
| | mean±sd | - | - | - | - | - | - | - | - | - | - | - | - | 0.054±0.024 |

**Table S2**. Isolate genomes used in metagenomic read recruitment analyses and clade identities.

| Strain Name | Organism | Clade | Accession |
|---|---|---|---|
| MED4 | Prochlorococcus sp. | HL_I | GCF_000011465.1 |
| MIT9515 | Prochlorococcus sp. | HL_I | GCF_000015665.1 |
| AS9601 | Prochlorococcus sp. | HL_II | GCF_000015645.1 |
| MIT9301 | Prochlorococcus sp. | HL_II | GCF_000015965.1 |
| MIT9215 | Prochlorococcus sp. | HL_II | GCF_000018065.1 |
| MIT9202 | Prochlorococcus sp. | HL_II | GCF_000158595.1 |
| MIT9107 | Prochlorococcus sp. | HL_II | GCF_000759855.1 |
| GP2 | Prochlorococcus sp. | HL_II | GCF_000759885.1 |
| MIT9201 | Prochlorococcus sp. | HL_II | GCF_000759955.1 |
| MIT9302 | Prochlorococcus sp. | HL_II | GCF_000759975.1 |
| MIT9311 | Prochlorococcus sp. | HL_II | GCF_000760015.1 |
| MIT9314 | Prochlorococcus sp. | HL_II | GCF_000760035.1 |
| MIT9321 | Prochlorococcus sp. | HL_II | GCF_000760055.1 |
| SB | Prochlorococcus sp. | HL_II | GCF_000760115.1 |
| NATL2A | Prochlorococcus sp. | LL_I | GCF_000012465.1 |
| NATL1A | Prochlorococcus sp. | LL_I | GCF_000015685.1 |
| PAC1 | Prochlorococcus sp. | LL_I | GCF_000760235.1 |
| CCMP1375 | Prochlorococcus sp. | LL_II | GCF_000007925.1 |
| MIT9211 | Prochlorococcus sp. | LL_III | GCF_000018585.1 |
| MIT9303 | Prochlorococcus sp. | LL_IV | GCF_000015705.1 |
| MIT9313 | Prochlorococcus sp. | LL_IV | GCF_000011485.1 |
| MITS9509 | Synechococcus sp. | CRD1 | GCF_001631935.1 |
| MITS9508 | Synechococcus sp. | CRD1 | GCF_001632165.1 |
| GEYO | Synechococcus sp. | CRD1 | GCF_900473955.1 |
| UW179A | Synechococcus sp. | CRD1 | GCF_900473965.1 |
| CC9311 | Synechococcus sp. | I | GCF_000014585.1 |
| WH8016 | Synechococcus sp. | I | GCF_000230675.1 |
| WH8020 | Synechococcus sp. | I | GCF_001040845.1 |
| UW179B | Synechococcus sp. | I | GCF_900474245.1 |
| CC9605 | Synechococcus sp. | II | GCF_000012625.1 |
| WH8109 | Synechococcus sp. | II | GCF_000161795.2 |
| N32 | Synechococcus sp. | II | GCF_900473895.1 |
| N5 | Synechococcus sp. | II | GCF_900473925.1 |
| N26 | Synechococcus sp. | II | GCF_900473975.1 |
| N19 | Synechococcus sp. | II | GCF_900474045.1 |
| UW86 | Synechococcus sp. | II | GCF_900474085.1 |
| WH8102 | Synechococcus sp. | III | GCF_000195975.1 |
| CC9902 | Synechococcus sp. | IV | GCF_000012505.1 |
| BL107 | Synechococcus sp. | IV | GCF_000153805.1 |
| RS9916 | Synechococcus sp. | IX | GCF_000153825.1 |
| CC9616 | Synechococcus sp. | UC-A | GCF_000515235.1 |
| KORDI100 | Synechococcus sp. | UC-A | GCF_000737535.1 |
| WH7803 | Synechococcus sp. | V | GCF_000063505.1 |
| WH7805 | Synechococcus sp. | VI | GCF_000153285.1 |
| RS9917 | Synechococcus sp. | VIII | GCF_000153065.1 |
| KORDI49 | Synechococcus sp. | WPC1 | GCF_000737575.1 |
| KORDI52 | Synechococcus sp. | WPC2 | GCF_000737595.1 |
| UW106 | Synechococcus sp. | XV | GCF_900474015.1 |
| UW69 | Synechococcus sp. | XV | GCF_900474185.1 |
| UW105 | Synechococcus sp. | XVI | GCF_900473935.1 |
| UW140 | Synechococcus sp. | XVI | GCF_900474295.1 |
| PCC6307 | Cyanobium gracile | SC_5.2 | GCF_000316515.1 |
| PCC7001 | Cyanobium sp. | SC_5.2 | GCF_000155635.1 |
| CB0101 | Synechococcus sp. | SC_5.2 | GCF_000179235.2 |
| RCC307 | Synechococcus sp. | SC_5.3 | GCF_000063525.1 |
| HIMB2401 | Synechococcus sp. | II | JBDJKK000000000 |
| PCC7421 | Gloeobacter violaceus | Outgroup | GCF_000011385.1 |

**Table S3.** Phytoplankton pigments across stations. Abbreviations: Tchla: Total Chlorophyll a; Fuco: Fucoxanthin, Peri: Peridinin; Allo: Alloxanthin; Prasino: Prasinoxanthin; Hex: 19′-hexanoyloxyfucoxanthin; But: 19′-butanoyloxyfucoxanthin; Zea: zeaxanthin; DVchl: divinyl chlorophyll *a*

| | HP1 | SB | SR8 | SR2 | STO1 | ALOHA |
|---|---|---|---|---|---|---|
| n | 10 | 3 | 10 | 10 | 10 | 25 |
| **But:Tchla** | 0.02±0.01 | 0.03±0.02 | 0.02±0.01 | 0.02±0.01 | 0.03±0.02 | 0.05±0.02 |
| **Hex:Tchla** | 0.04±0.01 | 0.04±0.01 | 0.04±0.01 | 0.04±0.03 | 0.07±0.04 | 0.13±0.03 |
| **Allo:Tchla** | 0.05±0.02 | 0.02±0.01 | 0.04±0.01 | 0.03±0.01 | 0.02±0.01 | - |
| **Fuco:Tchla** | 0.16±0.08 | 0.19±0.08 | 0.16±0.07 | 0.13±0.08 | 0.08±0.07 | 0.05±0.02 |
| **Lut:Tchla** | 0.01±0.01 | 0.01±0 | 0.01±0.01 | 0.01±0.01 | 0.01±0.01 | 0.01±0.01 |
| **Peri:Tchla** | 0.04±0.02 | 0.03±0.01 | 0.04±0.02 | 0.04±0.02 | 0.02±0.01 | 0.02±0.01 |
| **Prasino:Tchla** | 0.04±0.01 | 0.04±0.01 | 0.04±0.02 | 0.03±0.01 | 0.03±0.02 | - |
| **Zea:Tchla** | 0.18±0.11 | 0.19±0.07 | 0.18±0.11 | 0.17±0.09 | 0.31±0.13 | 0.64±0.18 |
| **DVa:Tchla** | 0.1±0.06 | 0.09±0.02 | 0.09±0.03 | 0.16±0.11 | 0.27±0.19 | 0.61±0.1 |

**Table S4.** Differences in biogeochemical parameters among community types.

| | | Nearshore | Transition | Offshore |
|---|---|---|---|---|
| | n | 229 | 85 | 52 |
| Seawater temp. (ºC) | mean±sd | 26.2±1.8 | 26.1±1.4 | 25.4±1.3 |
| pH | mean±sd | 7.9±0.2 | 7.9±0.2 | 8.0±0.2 |
| Salinity | mean±sd | 34.2±1.3 | 34.5±1.1 | 34.7±0.9 |
| Chlorophyll a (µg $L^{-1}$) | mean±sd | 1.7±2 | 0.8±0.5 | 0.4±0.3 |
| | n | 228 | 85 | 52 |
| Photosynthetic picoeukaryotes (cells $mL^{-1}$)*$10^4$ | mean±sd | 1.75±1.03 | 0.81±0.43 | 0.32±0.14 |
| Heterotrophic bacteria (cells $mL^{-1}$) *$10^6$ | mean±sd | 1.53±0.59 | 0.86±0.23 | 0.66±0.11 |
| Synechococcus (cells $mL^{-1}$) *$10^4$ | mean±sd | 20.78±14.42 | 5.94±4.43 | 1.96±1.69 |
| Prochlorococcus (cells $mL^{-1}$) *$10^4$ | mean±sd | 0.64±0.91 | 4.3±3.74 | 18.48±6.5 |
| | n | 107 | 33 | 27 |
| Phosphate (µM) | mean±sd | 0.11±0.07 | 0.08±0.03 | 0.08±0.03 |
| Silicate (µM) | mean±sd | 12.62±21.08 | 2.49±2.03 | 1.55±1.37 |
| Nitrate+Nitrite (µM) | mean±sd | 0.29±0.29 | 0.27±0.23 | 0.12±0.12 |
| | n | 41 | 10 | 3 |
| Ammonia (µM) | mean±sd | 0.27±0.36 | 0.12±0.16 | 0.04±0.02 |

**Table S5.** Alpha diversity metrics (Shannon's and ASV richness) across community types and across community types and seasons.

| Shannon's | Community Type | Estimates | Standard Errors | p-values |
|---|---|---|---|---|
| | Nearshore | 1.32 | 0.00 | 0 |
| | Transition | 1.68 | 0.01 | 0 |
| | Offshore | 1.45 | 0.03 | 0 |
| **ASV Richness** | **Community Type** | **Estimates** | **Standard Errors** | **p-values** |
| | Nearshore | 54 | 1 | 0 |
| | Transition | 47 | 2 | 0 |
| | Offshore | 44 | 2 | 0 |

**Table S6**. Significant differences in abundance across community types for phytoplankton genera as measured by ANCOM-BC2. FDR-corrected q-values are reported.

| | Offshore vs Nearshore | | | Transition vs Nearshore | | | Offshore vs Transition | | |
|---|---|---|---|---|---|---|---|---|---|
| Genus | log-fold change | q-val | sensitivity test passed | log-fold change | q-val | sensitivity test passed | log-fold change | q-val | sensitivity test passed |
| Micromonas | 0.164 | 0.543 | TRUE | -0.743 | 0.000 | TRUE | -0.907 | 0.001 | TRUE |
| unclassified Mamiellaceae* | 0.781 | 0.000 | TRUE | -0.399 | 0.035 | TRUE | -1.180 | 0.000 | TRUE |
| Bathycoccus | -0.910 | 0.000 | TRUE | -1.258 | 0.000 | TRUE | -0.348 | 0.164 | FALSE |
| Ostreococcus | 1.005 | 0.000 | TRUE | -0.407 | 0.045 | FALSE | -1.412 | 0.000 | TRUE |
| unclassified Bathycoccaceae* | 0.481 | 0.042 | FALSE | -0.862 | 0.000 | TRUE | -1.343 | 0.000 | TRUE |
| Mamiella | 0.730 | 0.000 | TRUE | -0.033 | 1.000 | TRUE | -0.762 | 0.002 | TRUE |
| unclassified Cryptomonadales* | -0.098 | 0.908 | FALSE | -0.939 | 0.000 | TRUE | -0.841 | 0.001 | TRUE |
| Proteomonas | 0.492 | 0.019 | TRUE | -0.433 | 0.024 | TRUE | -0.925 | 0.000 | TRUE |
| Teleaulax | 0.081 | 0.972 | TRUE | -1.163 | 0.000 | TRUE | -1.244 | 0.000 | TRUE |
| Tetraselmis | 0.097 | 1.000 | FALSE | -1.127 | 0.000 | TRUE | -1.223 | 0.000 | TRUE |
| Chloropicaceae* | -1.377 | 0.000 | TRUE | -2.474 | 0.000 | TRUE | -1.098 | 0.000 | TRUE |
| Ochromonas | -0.587 | 0.005 | TRUE | -0.043 | 1.000 | TRUE | 0.544 | 0.037 | TRUE |
| Prasinoderma | 0.460 | 0.033 | TRUE | -0.335 | 0.092 | FALSE | -0.795 | 0.002 | TRUE |
| Atelocyanobacterium (UCYN-A) | 0.220 | 0.367 | FALSE | 1.388 | 0.000 | TRUE | 1.168 | 0.000 | TRUE |
| Pyramimonas | -0.358 | 0.102 | TRUE | -1.360 | 0.000 | TRUE | -1.003 | 0.000 | TRUE |
| Cyanobium | -0.853 | 0.000 | TRUE | -2.538 | 0.000 | TRUE | -1.685 | 0.000 | TRUE |
| Prochlorococcus | 2.240 | 0.000 | TRUE | 4.366 | 0.000 | TRUE | 2.126 | 0.000 | TRUE |
| Synechococcus | -0.475 | 0.012 | TRUE | -1.512 | 0.000 | TRUE | -1.037 | 0.000 | TRUE |
| Eutreptiella | 0.483 | 0.056 | FALSE | -0.204 | 0.502 | FALSE | -0.686 | 0.029 | TRUE |
| unclassified Pelagomonadaceae* | 0.308 | 0.178 | FALSE | 0.756 | 0.001 | TRUE | 0.448 | 0.178 | FALSE |
| Sarcinochrysidaceae* | 1.381 | 0.000 | TRUE | 1.859 | 0.000 | TRUE | 0.478 | 0.062 | FALSE |
| Chaetoceros | -0.583 | 0.005 | TRUE | -1.812 | 0.000 | TRUE | -1.229 | 0.000 | TRUE |
| unclassified Polar-centric-Mediophyceae | -0.217 | 0.449 | TRUE | -2.035 | 0.000 | TRUE | -1.818 | 0.000 | TRUE |
| Phaeocystis | 0.261 | 0.229 | TRUE | 1.032 | 0.000 | TRUE | 0.771 | 0.002 | TRUE |
| Chrysochromulinaceae* | -1.192 | 0.000 | FALSE | -0.105 | 0.938 | FALSE | 1.087 | 0.000 | TRUE |
| Chrysochromulina | 0.243 | 0.276 | FALSE | -0.546 | 0.010 | TRUE | -0.788 | 0.002 | TRUE |
| Braarudosphaera | 0.158 | 0.583 | FALSE | 1.195 | 0.000 | TRUE | 1.036 | 0.000 | TRUE |
| unclassified Noelaerhabdaceae* | -0.455 | 0.027 | TRUE | -0.664 | 0.001 | TRUE | -0.209 | 0.503 | FALSE |

**Table S7.** Seasonal differences in biogeochemical parameters across community types. Comparisons of biogeochemical parameters between winter and summer seasons using one-way ANOVA. Significance (uncorrected p-values) and F-statistic are shown. Degrees of freedom equal to one. The pound sign (#) denotes a log transformation to improve normality, while a carrot (^) denotes a log transformation plus 1.

| | | Nearshore Summer | Nearshore Winter | Transition Summer | Transition Winter | Offshore Summer | Offshore Winter | Nearshore Summer-Winter | | Transition Summer-Winter | | Offshore Summer-Winter | |
|---|---|---|---|---|---|---|---|---|---|---|---|---|---|
| | n | 51 | 61 | 27 | 13 | 14 | 18 | p | F | p | F | p | F |
| Seawater temp. (ºC) | mean±sd | 28.1±0.9 | 24.1±0.7 | 27.4±0.5 | 24.2±0.4 | 26.7±0.5 | 24.2±0.4 | <0.001 | 697.6 | <0.001 | 366.1 | <0.001 | 180.1 |
| pH | mean±sd | 7.9±0.2 | 7.9±0.2 | 8.0±0.2 | 8.0±0.2 | 8.1±0.2 | 8.0±0.2 | 0.789 | 0.006 | 0.48 | 0.509 | 0.211 | 1.632 |
| Salinity | mean±sd | 33.7±1.2 | 33.8±1.3 | 33.9±0.9 | 34.6±0.9 | 34.3±0.8 | 34.6±0.8 | 0.572 | 0.321 | 0.022 | 5.743 | 0.234 | 1.476 |
| Chlorophyll a (μg $L^{-1}$)# | mean±sd | 1.5±1.0 | 2.3±3.4 | 0.8±0.5 | 0.7±0.4 | 0.4±0.2 | 0.4±0.3 | 0.031 | 4.774 | 0.43 | 0.637 | 0.897 | 0.017 |
| | n | 51 | 61 | 27 | 13 | 14 | 18 | | | | | | |
| Photosynthetic picoeukaryotes (cells $mL^{-1}$)*$10^{4}$# | mean±sd | 2.14±1.31 | 1.60±1.23 | 1.01±0.51 | 0.51±0.40 | 0.29±0.11 | 0.39±0.18 | 0.004 | 8.425 | <0.001 | 20.63 | 0.08 | 3.279 |
| Heterotrophic bacteria (cells $mL^{-1}$) *$10^{6}$# | mean±sd | 1.81±0.64 | 1.39±0.47 | 0.99±0.23 | 0.73±0.22 | 0.72±0.92 | 0.63±0.13 | <0.001 | 18.81 | 0.004 | 15.31 | 0.019 | 6.11 |
| Synechococcus (cells $mL^{-1}$) *$10^{4}$# | mean±sd | 30.09±15.04 | 14.99±14.77 | 8.88±4.65 | 1.75±1.79 | 3.56±1.92 | 0.91±0.61 | <0.001 | 38.29 | <0.001 | 66.98 | <0.001 | 53.09 |
| Prochlorococcus (cells $mL^{-1}$) *$10^{4}$^ | mean±sd | 0.21±0.43 | 0.90±0.78 | 3.22±3.23 | 3.98±4.44 | 16.55±4.43 | 18.20±7.24 | <0.001 | 109.7 | 0.233 | 1.471 | 0.814 | 0.056 |
| | n | 20 | 35 | 10 | 6 | 7 | 11 | | | | | | |
| Phosphate (μM)# | mean±sd | 0.15±0.07 | 0.1±0.06 | 0.11±0.04 | 0.07±0.02 | 0.09±0.03 | 0.09±0.03 | 0.004 | 8.915 | 0.036 | 5.399 | 0.747 | 0.108 |
| Silicate (μM)# | mean±sd | 16.48±24.81 | 10.16±19 | 3.14±2.99 | 2.42±1.29 | 2.38±2.62 | 1.3±0.15 | 0.04 | 4.448 | 0.75 | 0.106 | 0.172 | 2.041 |
| Nitrate+Nitrite (μM)# | mean±sd | 0.33±0.43 | 0.27±0.23 | 0.3±0.32 | 0.18±0.06 | 0.11±0.06 | 0.1±0.07 | 0.637 | 0.225 | 0.961 | 0.003 | 0.714 | 0.139 |
| | n | 7 | 16 | 3 | 1 | 1 | 1 | | | | | | |
| Ammonia (μM)# | mean±sd | 0.4±0.49 | 0.26±0.38 | 0.06±0.02 | | | | 0.226 | 1.556 | | | | |
| | n | 9 | 9 | | | | | | | | | | |
| Max rainfall (mm)^ | mean±sd | 1.52±2.29 | 10.92±14.48 | | | | | 0.036 | 5.256 | | | | |
| Max wind speed ($ms^{-1}$) | mean±sd | 9.13±3.66 | 23.01±2.48 | | | | | 0.084 | 3.388 | | | | |
| Average wind direction (°)^ | mean±sd | 76.48±26.72 | 144.77±37.36 | | | | | <0.001 | 20.73 | | | | |

**Table S8.** Phytoplankton genera with significant seasonality as defined by Lomb-Scargle Periodicity test. FDR-corrected q-values reported.

| Community type | Genus | Lomb-Scargle Periodicity (LSP) q-val |
|---|---|---|
| Nearshore | unclassified Pycnococcaceae | <0.001 |
| Nearshore | *Prochlorococcus* | <0.001 |
| Nearshore | *Mesopedinella* | 0.007 |
| Nearshore | *Chaetoceros* | <0.001 |
| Nearshore | unclassified Chloropicaceae | 0.002 |
| Nearshore | *Teleaulax* | 0.003 |
| Nearshore | unclassified Mamiellaceae | <0.001 |
| Nearshore | *Micromonas* | 0.001 |
| Nearshore | *Isochrysis* | <0.001 |
| Nearshore | *Partenskyella* | <0.001 |
| Nearshore | unclassified Polar-centric Mediophyceae | <0.001 |
| Nearshore | *Helicopedinella* | 0.003 |
| Nearshore | *Eutreptiella* | <0.001 |
| Nearshore | *Synechococcus* | <0.001 |
| Nearshore | *Chrysochromulina* | <0.001 |
| Nearshore | *Braarudosphaera* | <0.001 |
| Nearshore | *Cyanobium* | <0.001 |
| Nearshore | *Mamiella* | 0.001 |
| Transition | *Prasinoderma* | <0.001 |
| Transition | *Pyramimonas* | 0.004 |
| Transition | *Synechococcus* | <0.001 |
| Transition | *Braarudosphaera* | 0.003 |
| Transition | *Cyanobium* | <0.001 |
| Offshore | *Atelocyanobacterium* | <0.001 |
| Offshore | *Micromonas* | <0.001 |
| Offshore | *Crocosphaera* | <0.001 |
| Offshore | unclassified Pelagomonadaceae | <0.001 |
| Offshore | *Synechococcus* | 0.005 |
| Offshore | *Braarudosphaera* | <0.001 |

**Table S9.** Alpha diversity metrics (Shannon's and ASV richness) across community types and seasons.

| Shannon's | Community Type | Season | Estimates | Standard Errors | p-values |
|---|---|---|---|---|---|
| | Nearshore | Summer | 1.15 | 0 | 0 |
| | | Winter | 1.66 | 0.01 | 0 |
| | Transition | Summer | 1.47 | 0.02 | 0 |
| | | Winter | 2.39 | 0.04 | 0 |
| | Offshore | Summer | 1.38 | 0.01 | 0 |
| | | Winter | 1.73 | 0.02 | 0 |
| **ASV Richness** | **Community Type** | **Season** | **Estimates** | **Standard Errors** | **p-values** |
| | Nearshore | Summer | 45 | 1 | 0 |
| | | Winter | 54 | 2 | 0 |
| | Transition | Summer | 43 | 2 | 0 |
| | | Winter | 50 | 3 | 0.047 |
| | Offshore | Summer | 35 | 2 | 0 |
| | | Winter | 47 | 2 | 0 |
